# Nicotinamide riboside alleviates Parkinson’s disease symptoms but downregulates dopamine metabolism upon lactacystin-induced proteostasis failure

**DOI:** 10.1101/2021.03.12.435062

**Authors:** Giorgio Turconi, Farhan Alam, Tanima SenGupta, Sini Pirnes-Karhu, Soophie Olfat, Mark S. Schmidt, Kärt Mätlik, Ana Montaño-Rodriguez, Vladimir Heiskanen, Petteri T. Piepponen, Charles Brenner, Carina I. Holmberg, Hilde Nilsen, Jaan-Olle Andressoo, Eija Pirinen

## Abstract

Activation of mitochondrial metabolism and proteostasis with the NAD^+^ precursor nicotinamide riboside (NR) has emerged as a potential therapeutic approach for neurodegenerative disorders including Parkinson’s disease (PD). However, despite recently started clinical trials, studies on NR in animal models of PD are scarce. In this study, we investigated the effect of NR in multiple models of PD. In transgenic *C. elegans* overexpressing α-synuclein, a protein of which aggregation is believed to promote PD, NR rescued PD-like phenotypes including mitochondrial dysfunction and motility defects, decreased oxidative stress, and age-related dopamine (DA) neuron loss. We found that NR eased symptoms of disease by activating the mitochondrial unfolded protein response (UPR^mt^) via the transcription factor *atfs-1*. Similarly, in a proteasome inhibitor, lactacystin, -induced mouse model of PD, NR rescued mitochondrial dysfunction and behavioural deficits caused by lactacystin lesion. However, long-term NR supplementation, in conjunction with proteasome inhibition, resulted in decreased DA levels in both the lesioned and unlesioned sides of the substantia nigra with concomitant downregulation of key genes in DA metabolism. Our results suggest specific endpoints that should be monitored in ongoing NR clinical trials.

## Introduction

Parkinson’s disease (PD) is a neurodegenerative disorder affecting more than 10 million people worldwide (1). Common symptoms of PD are motor abnormalities including bradykinesia, resting tremor, and muscle rigidity caused by progressive degeneration of dopamine (DA) neurons in the midbrain substantia nigra pars compacta (2). Currently, there is no disease modifying treatment for PD.

Substantial evidence suggests that defects in proteostasis and mitochondrial function are the main drivers of PD (3) and that an intimate interplay between these two processes exists. Failure in the ubiquitin-proteasome system, including a reduction in proteasome function and the loss of 26S/20S proteasome subunits, have been reported in the substantia nigra of PD patients (4, 5). In PD, mitochondrial dysfunction typically manifests itself as impaired mitochondrial respiratory capacity and quality control, which have been suggested to be caused by an interplay between genetic (6) and environmental factors (7, 8). Mitochondrial dysfunction may promote the formation of protein aggregations by inducing oxidative stress or disturbing the proteasome function via ATP deficit. Impaired proteostasis subsequently leads to the appearance of Lewy bodies, resulting from the overproduction and misfolding of α-synuclein (α-syn) within the soma and neurites of neurons (9). On the other hand, α-syn aggregates disrupt the activity of the mitochondrial electron transport chain (ETC) complexes either directly or indirectly by suppressing mitochondrial proteases and chaperones (10, 11). These proteases and chaperons control the turnover of mitochondrial matrix proteins, including components of ETC, and play an important role in the mitochondrial unfolded protein response (UPR^mt^), a signalling pathway that confers mitochondrial proteostasis in response to stress. The vicious cycle of mitochondrial dysfunction resulting in Lewy body formation and overproduction of α-syn further disturbing mitochondrial function provides a rationale for enhancing mitochondrial function as treatment for PD.

Recently, activation of mitochondrial capacity and quality control, based mainly on lifestyle intervention studies, has emerged as a promising option for the treatment of neurodegenerative disorders (12). Currently, there are few pharmacological approaches available to promote mitochondrial function, with one of the most promising compounds being nicotinamide riboside (NR), an NAD^+^ precursor vitamin B3 (13). In the cell, NR is used for the biosynthesis of nicotinamide adenine dinucleotide (NAD^+^), the central co-enzyme of mitochondrial energy generation. Mounting evidence suggests that the maintenance of proper NAD^+^ homeostasis is essential for DA neurons that are special types of neurons differing from most of the other neurons in the central nervous system in several ways, including greater energy demand, increased mitochondrial activity, reactive oxygen species release, and altered Ca^+^ homeostasis (14, 15). Interestingly, disrupted NAD^+^ metabolism has been reported in PD (16–18). In preclinical studies, NR has shown great promise regarding the treatment of neurodegenerative diseases by rescuing NAD^+^ homeostasis and mitochondrial oxidative capacity, and by decreasing the formation of amyloid-beta and superoxide dismutase −1 protein aggregations in mouse models of Alzheimer’s disease and amyotrophic lateral sclerosis, respectively (19, 20). In addition, NR has been observed to rescue mitochondrial defects in PD patient induced pluripotent stem cell-derived DA neurons and to alleviate motor deficits in a fly model of PD (17). In previous studies, the beneficial effect of NR on mitochondrial and protein homeostasis has been suggested to be mediated via an activation of UPR^mt^ and the subsequent restoration of mitochondrial proteostasis (19, 20). Encouraged by the success of NR in various animal models of human disease, two clinical trials with NR are ongoing in PD patients at the time of writing (NCT03568968, NCT04044131), and one clinical Phase I trial with a short-term NR supplementation of four weeks is already completed (NCT03816020). However, to evaluate efficacy and safety, the analysis of new treatment options in PD animal models is important. Because an animal model that phenocopy the slow progression of PD in humans is not available, the only route forward in establishing treatment safety and efficacy is a careful analysis of various available animal models.

In this study, we determine the effect of NR on PD-like phenotypes in transgenic *C. elegans* strains overexpressing α-syn in either the body wall muscle cells or in DA neurons, as well as in a 26S/20S proteasome inhibitor lactacystin mouse model of PD. We find that NR attenuates PD associated disease symptoms both in *C. elegans* and in mice, but the primary endpoint - a rescue of nigrostriatal dopamine levels and function - is not reached. Instead, NR disturbs DA metabolism upon drug-induced proteostasis failure in mice. Our results suggest specific endpoints to be added for clinical monitoring and call for further research in various PD models.

## Results

### NR supplementation corrects disease-like phenotypes in *C. elegans* models of PD

First, we tested a *C. elegans* model of PD overexpressing human wild-type α-syn in the body wall muscle cells (muscle α-syn) (21). To determine the effect of NR on α-syn overexpression induced pathogenic processes, muscle α-syn and control animals were supplemented with NR and vehicle from the L1 larval stage. Basal and maximal respiration rates were significantly decreased in muscle α-syn *C. elegans* from day 2 of adult age onwards, confirming that this strain exhibited mitochondrial dysfunction (Figure 1A). However, the amount of mtDNA was significantly higher in the muscle α-syn strain, suggesting a compensatory increase in mitochondrial content to rescue mitochondrial oxidative capacity (Supplemental Figure 1A). Supplementation with the NAD^+^ precursor vitamin B3, NR, resulted in increased NAD^+^ levels in both strains, confirming that NR was metabolized as expected (Figure 1B). There was no change in pharyngeal pumping activity (*i.e.* food intake) between the strains during early adult age, but the muscle α-syn strain displayed impaired food intake during late adult age, on day 6 (Supplemental Figure 1B). However, this was rescued by NR. In fact, NR supplementation improved pharyngeal pumping activity in both control and muscle α-syn over-expressing *C. elegans* strains at the early and late adult ages (Supplemental Figure 1B). Notably, NR supplementation effectively rescued both basal and maximal respiration rates at days 2 and day 4 of adult age (Figure 1A) in muscle α-syn strain, but did not increase the mtDNA level, similarly to the control strain (Supplemental Figure 1A). In line with the improvement in mitochondrial function, NR rescued motility defects in muscle α-syn strain from day 4 until day 8 of adult age (Figure 1C). Altogether, these results suggest that NR-mediated mitochondrial activation decreases α-syn overexpression-driven decline in motility and mitochondrial function within the body wall muscle.

**Figure 1:**
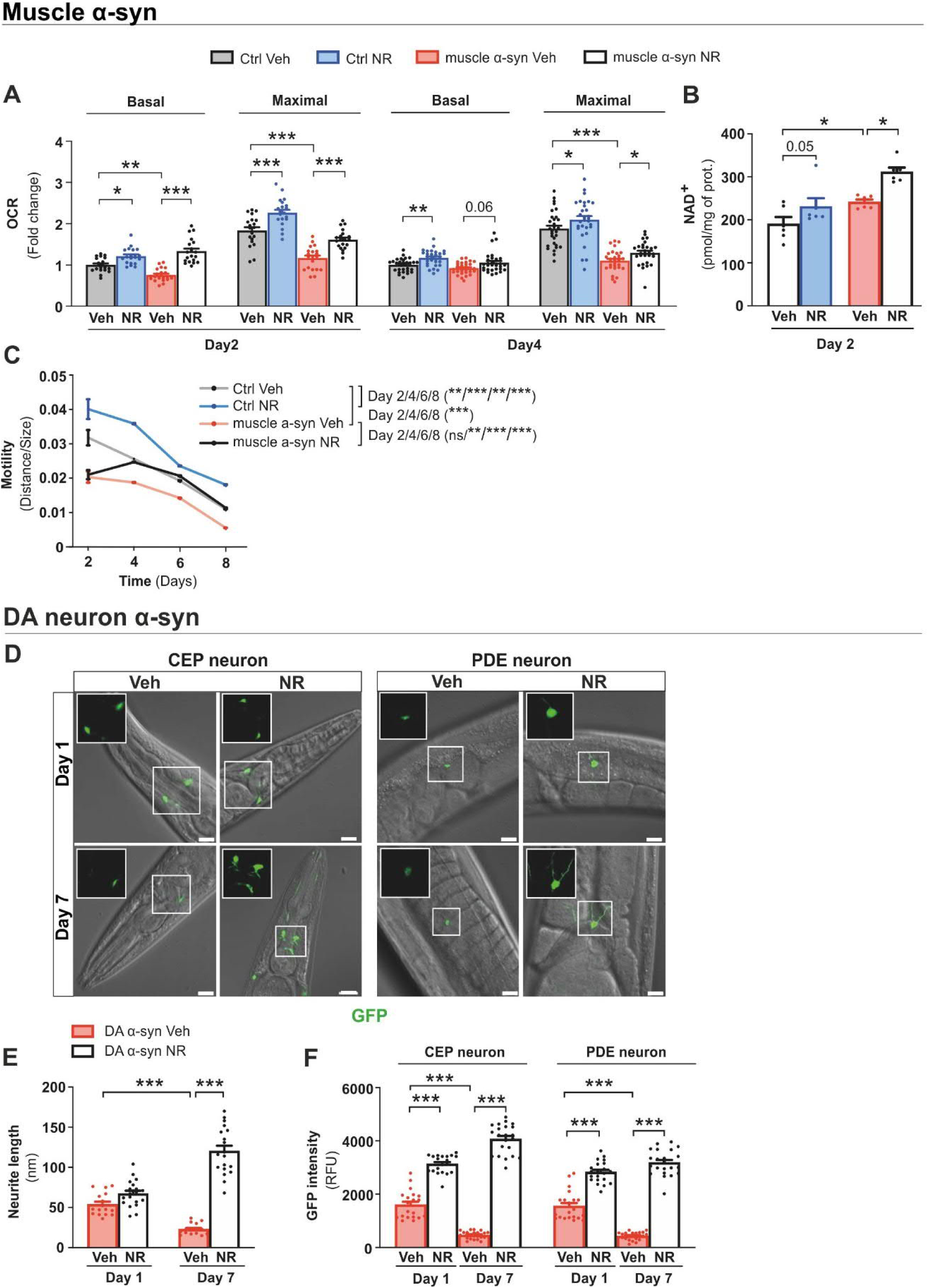
NR alleviates PD-like phenotypes in α-syn overexpressing *C. elegans* models, **(A)** Basal and maximal mitochondrial respiration after carbonyl cyanide-4-(trifluoromethoxy)phenylhydrazone (15 µM) injection measured in the control and muscle α-syn strains after vehicle (Veh) or NR supplementation at days 2 and 4 of adult age. Each data point represents the mean of 2 independent experiments in total with 20 wells per group, with each group having 5 to 8 animals per well. (**B**) Total NAD^+^ levels in the vehicle and NR supplemented control and muscle α-syn strains at day 2 of adult age measured by modified colorimetric enzymatic method (n=6 biologically independent samples per group and two independent experiments). (**C)** Motility (distance travelled normalized to the size of the animals) in the control and muscle α-syn strains with vehicle or NR supplementation at days 4, 6, and 8 of adult age. Each data point represents the mean of 2 independent experiments with 59 to 99 animals per group. (**D)** Confocal images of CEP and PDE DA neurons and neurite extensions in the DA α-syn strain after vehicle or NR supplementation at days 1 and 7 of adult age. Inserts show CEP and PDE neurites with 63X magnification. Scale bar, 5 µm. (**E)** Neurite length in the DA α-syn strain after vehicle or NR supplementation at days 1 and 7 of adult age and (**F**) quantification of GFP signal intensity in CEP and PDE DA neurons in the DA α-syn strain using ImageJ software (n=20 to 23 biologically independent samples per group and two independent experiments), analysed from the confocal images represented in panel (**D**) using ImageJ software (n=16 to 19 biologically independent samples per group and two independent experiments). All data are mean ± SEM \**P* < 0.05; \*\**P* ≤ 0.01; \*\*\**P* ≤ 0.001; ns, not significant. Statistical analysis was performed in GraphPad prism version 9.0.0. Overall differences between conditions were assessed in one-way ANOVA using an uncorrected version of Fisher’s LSD test. OCR, oxygen consumption rate; GFP, green fluorescent protein; DA, dopamine; CEP, cephalic; PDE, posterior deirid. See also: Supplemental Figure 1.

To analyse the effect of NR on DA neurons, we subsequently used another transgenic strain specifically co-expressing GFP with α-syn in DA neurons (DA α-syn). This *C. elegans* model demonstrates age-dependent dopaminergic neuron degeneration (22). DA neuron specific α-syn overexpression sufficiently compromised basal respiration in young DA α-syn animals (Supplemental Figure 1C). This mitochondrial bioenergetic defect was corrected by NR supplementation in young animals (Supplemental Figure 1C). Furthermore, NR provided a complete rescue of α-syn overexpression as well as aging-driven DA neurite (Figure 1D, E) and neuron loss (Figure 1D, F). NR also maintained normal feeding behaviour and muscle fitness upon aging in DA α-syn and control strains (Supplemental Figure 1D, E). Thus, NR protects from the α-syn overexpression-induced loss of DA neurites and neurons.

### NR mitigates disease phenotypes via an UPR^mt^ dependent mechanism in *C. elegans*

As NR has been previously shown to mediate its beneficial effect on neurodegeneration via the mitochondrial unfolded protein (UPR^mt^) pathway in *C. elegans* models of Alzheimer’s disease (20), we next investigated this adaptive transcriptional response pathway using the muscle α-syn strain. We observed a clear signature of repressed UPR^mt^ signalling in our PD model, since the mitochondrial protease genes *ymel-1, lonp-1,* and *clpP,* as well as the positive regulator of UPR^mt^ signalling, *ubl-5,* were significantly downregulated in muscle α-syn strain when compared to controls (Figure 2A). Notably, NR activated UPR^mt^ signalling by increasing the expression of *lonp-1*, *clpP*, and *ubl-5* in the muscle α-syn strain (Figure 2A). Therefore, these findings suggest that the suppression of UPR^mt^ signalling is one mechanism of α-syn toxicity, which is reversed upon NR supplementation.

**Figure 2:**
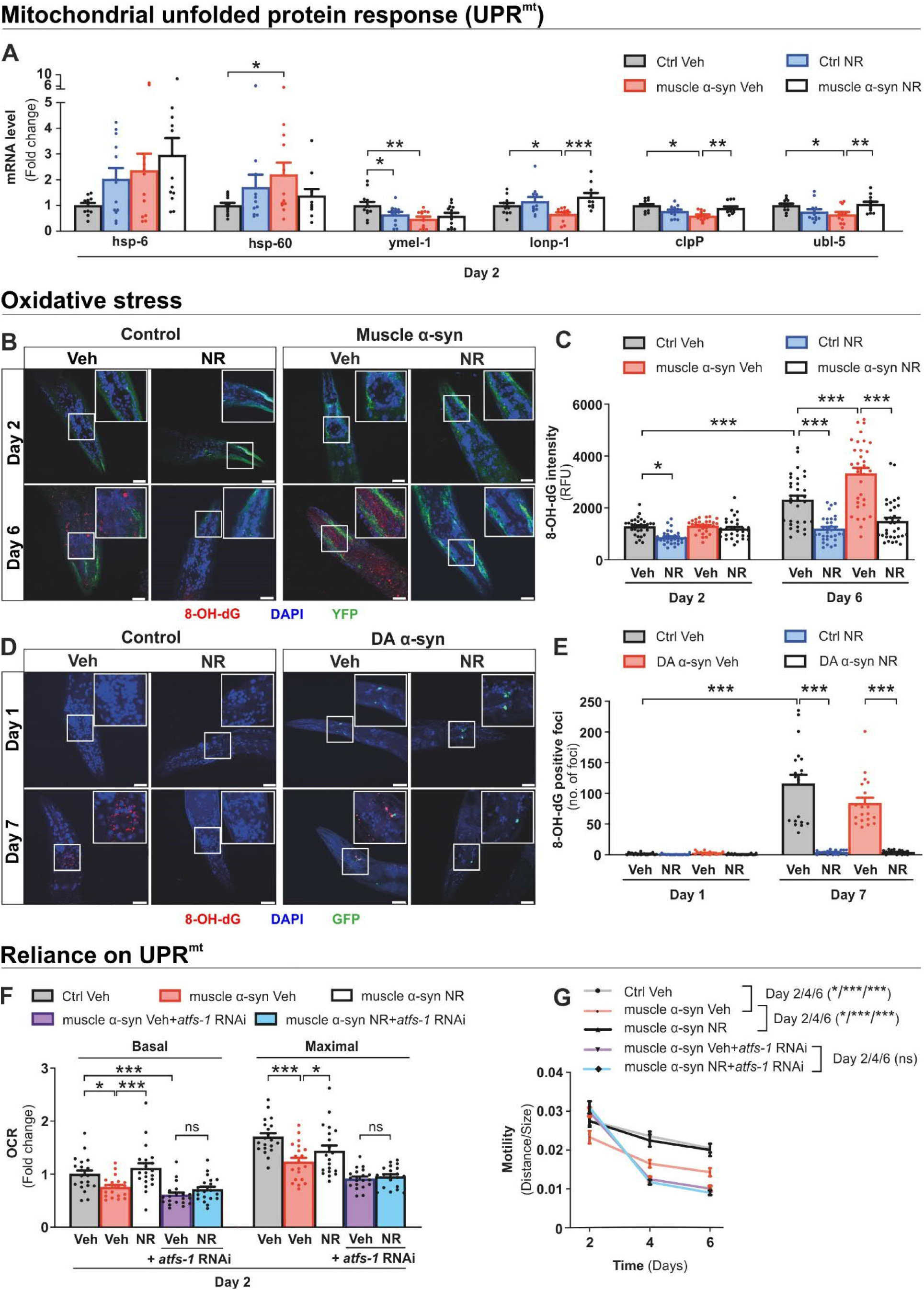
NR supplementation decreases oxidative stress and improves disease-like phenotypes via UPR^mt^. **(A)** Messenger RNA expression of genes involved in UPR^mt^ in the control and muscle α-syn *C. elegans* strains after vehicle (Veh) and NR supplementation at day 2 of adult age. Data represent the mean of 2 independent experiments with pools of approximately 1500 animals per group. (**B)** Representative confocal images with higher magnification inserts of immunofluorescence staining of 8-OH-dG (red), the oxidative stress marker, in the control and muscle α-syn strains after vehicle or NR supplementation at days 2 and 6 of adult age. Nuclei were stained with DAPI (blue) and muscle α-syn accumulations were visualized by imaging the fluorescence signal from α-syn-YFP, compared to YFP in the control strain. Scale bar, 20 µm. (**C)** Corresponding quantification of fluorescence intensity of 8-OH-dG staining in the control and muscle α-syn strains, analysed using ImageJ software (n=30 to 33 biologically independent samples per group and two independent experiments). (**D)** Representative confocal images with higher magnification inserts of immunofluorescence staining of 8-OH-dG (red), the oxidative stress marker, in the control, N2 wild type, and DA α-syn strains at days 1 and day 7 of adult age (n=19 biologically independent samples per group and two independent experiments). Scale bar, 20 µm. (**E)** Corresponding quantification of 8-OH-dG positive foci counted manually from the head region in the control and DA α-syn strains after vehicle or NR supplementation. (**F)** Basal and maximal mitochondrial respiration at day 2 of adult age in the vehicle treated control strain and muscle α-syn animals exposed to control or *atfs-1* RNAi. Data represent the mean of 2 independent experiments in total, each with 20 wells per group containing 5 to 8 animals per well. (**G)** Motility (distance travelled normalised to the size of the animals) in the vehicle treated control strain and muscle α-syn *C. elegans* fed with or without *atfs-1* RNAi at days 2, 4, and 6 of adult age. Each data point represents the mean of 2 independent experiments with 71 to 150 animals per group. All data are mean ± SEM. \**P* < 0.05; \*\**P* ≤ 0.01; \*\*\**P* ≤ 0.001; ns, not significant. Statistical analysis was performed in GraphPad Prism version 9.0.0. Overall differences between conditions were assessed with one-way ANOVA followed by an uncorrected version of Fisher’s LSD test. *Hsp6*, heat shock protein 6; *Hsp60*, heat shock protein 60; *ymel-1*, ATP-dependent zinc metalloprotease; *lonp-1*, Lon protease; *clpp-1,* ATP-dependent Clp protease proteolytic subunit 1; *ubl-5*, ubiquitin-like protein subunit 5; OCR, oxygen consumption rate; UPR^mt^, mitochondrial unfolded protein response.

Oxidative stress is strongly associated with UPR^mt^ signalling. An excessive production of reactive oxygen species (ROS) can stimulate UPR^mt^ in *C. elegans* (23) but, on the other hand, UPR^mt^ activation confers protection from oxidative stress via *atfs-1*, the key UPR^mt^ transcription factor, -mediated transcriptional response (24). Given that NR has been shown to trigger a strong burst of ROS in early adult age, yet has also been shown to stimulate a robust ROS defense in late adult age in *C. elegans* (25), we determined tissue oxidative stress immunohistochemically using an anti-8-OH-dG antibody. As expected, both muscle and DA α-syn *C. elegans* strains showed distinct anti-8-OH-dG foci upon aging (Figure 2B, D). Quantification of staining intensity clearly showed that NR did not elevate an accumulation of 8-OH-dG in young adults, but significantly decreased the levels of this widespread oxidative stress marker in both *C. elegans* strains at the later age of adulthood (Figure 2C, E). Thus, our results support that NR provides protection against oxidative stress induced upon aging. To understand whether UPR^mt^ was essential for NR-induced improvements in the α-syn overexpression PD model, we used genetic intervention by RNAi feeding to knock down *atfs-1*. The silencing of *atfs-1* abrogated the mitochondrial boosting effect of NR on day 2 of adult age in the muscle α-syn strain (Figure 2F). This was accompanied by the loss of NR-induced improvement in motility from day 4 of adulthood onwards in the muscle α-syn strain (Figure 2G). Overall, these results demonstrated that UPR^mt^ is one mechanism mediating the protective capacity of NR in the *C. elegans* model of PD.

### NR supplementation improves NAD^+^ metabolism in the brain in lactacystin mouse model of PD

Because NR supplementation improved disease related features in the *C. elegans* α-syn overexpression models, we subsequently studied the effect of NR in a mouse model of PD induced by the proteasome inhibitor lactacystin. We chose the lactacystin model over other toxin-based PD models because lactacystin recapitulates the general proteostasis failure by inhibiting 26S/20S function known to be reduced in substantia nigra in PD (26, 27) and leads to a reproducible lesion in the nigrostriatal DA system in mice (28–31) (see Study Design in Materials and Methods for further details). Mice were preconditioned with either vehicle or NR diets for 3 months prior to an induction of Parkinsonism through a unilateral stereotactic injection of lactacystin just above the substantia nigra (Figure 3A). Behavioural phenotype was studied 2 weeks before and 2, 6, and 10 weeks after the lactacystin injection (Figure 3A). Mice were sacrificed and dissected 12 weeks after the lactacystin injection. Given that there was no difference in food intake between vehicle and NR supplemented mice (Supplemental Figure 2A), NR intake was assumed to be similar in both groups. NR supplementation did not significantly affect body weight during the first 12 weeks of NR supplementation (Supplemental Figure 2B). However, NR-supplemented mice maintained their body weight more effectively after a supranigral injection of lactacystin, which caused a drop in body weight in both groups (Supplemental Figure 2B). Therefore, NR likely improved general well-being in lactacystin-injected PD mice.

**Figure 3.**
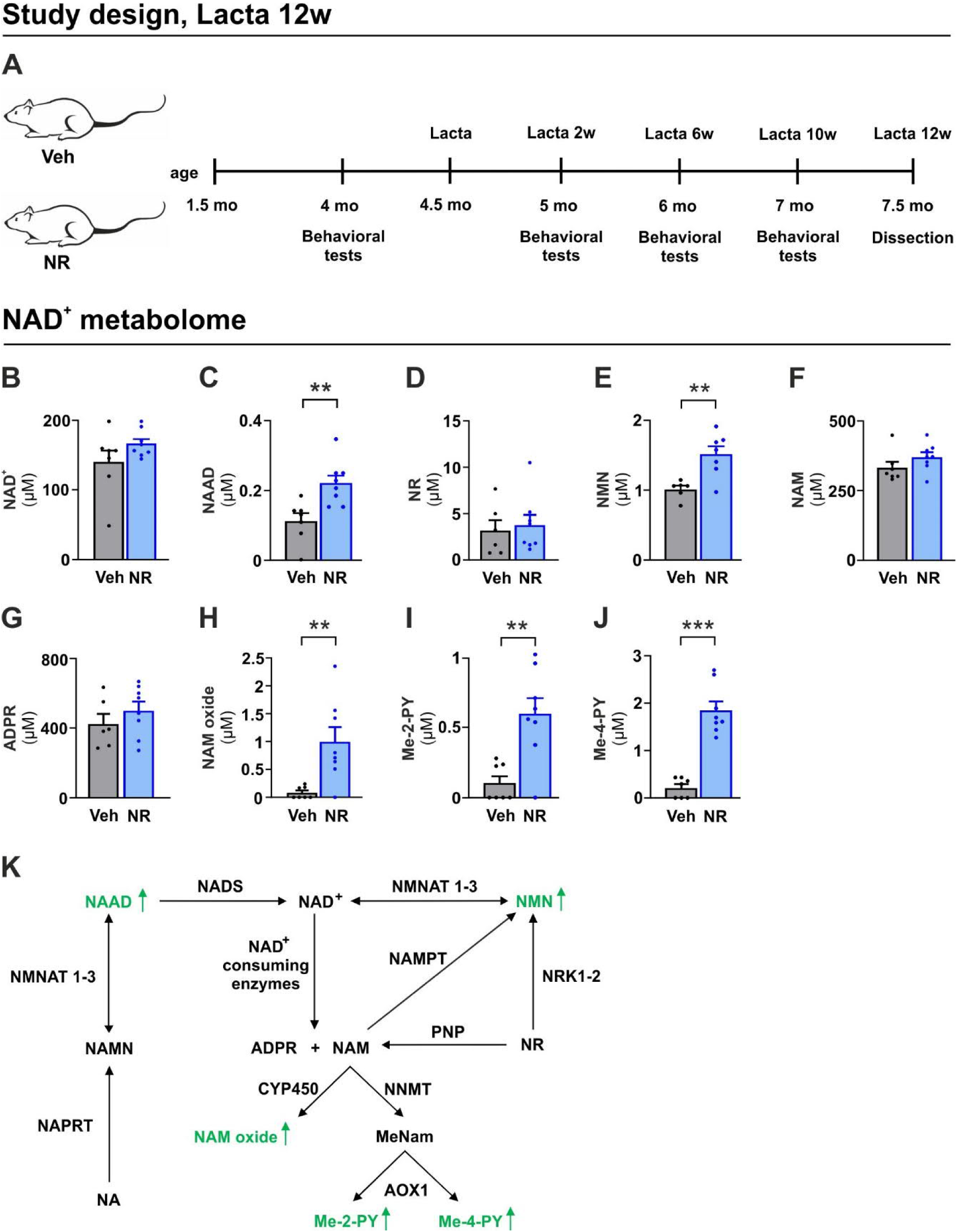
NR supplementation improves NAD^+^ metabolism in the brain. (**A**) Experimental design for a study in which PD-associated phenotypes of mice fed with either a vehicle (Veh) or NR-enriched diet (NR) were analyzed 12 weeks after nigral lactacystin lesion. Cerebellar content of (**B**) NAD^+^; (**C**) NAAD; (**D**) NR; (**E**) NMN; (**F**) NAM; (**G**) ADPR; (**H**) NAM oxide; (**I**) Me-2-PY; (**J**) Me-4-PY in vehicle and NR supplemented mice at the 12-week time point. Veh (n = 7) and NR (n = 8). (**K**) Schematic of the NAD^+^ metabolome. NAD metabolites that significantly increased upon NR supplementation are presented with the green font. Data are presented as mean ± SEM. **P ≤ 0.01, ***P ≤ 0.001. Statistical analysis was performed in GraphPad prism version 8.0.2. Overall differences between the two groups were assessed with Student’s t-test followed by Welch’s correction. ADPR, Adenosine diphosphate ribose; AOX1, aldehyde oxidase; CYP450, cytochrome P450; Lacta, lactacystin; Me-2-PY, N-methyl-2-pyridone-5-carboxamide; Me-4-PY, N-methyl-4-pyridone-5-carboxamide; MeNam, methylnicotinamide; NA, nicotinic acid; NAAD, nicotinic acid adenine dinucleotide; NAD^+^, nicotinamide adenine dinucleotide; NADS, nicotinamide adenine dinucleotide synthetase; NAM; nicotinamide; NAM oxide, nicotinamide N-oxide; NAMN, nicotinic acid mononucleotide; NAMPT, nicotinamide phosphoribosyltransferase; NAPRT, nicotinate phosphoribosyltransferase; NMN nicotinamide mononucleotide; NMNAT, nicotinamide mononucleotide adenylyltransferase; NNMT, nicotinamide N-methyltransferase; NR, nicotinamide riboside; NRK, nicotinamide riboside kinase; PNP, purine nucleoside phosphorylase; Veh, vehicle.

To study whether NR supplementation affected NAD^+^ metabolism in the brain, we performed a targeted liquid chromatography–mass spectrometry analysis of the cerebellum. The level of NAD^+^ did not differ between vehicle and NR supplemented mice (Figure 3B). However, we found elevated levels of nicotinic acid adenine dinucleotide (NAAD), a validated biomarker of increased NAD^+^ metabolism, upon NR administration (Figure 3C) (32). The level of NR remained unaltered (Figure 3D) while the level of nicotinamide mononucleotide (NMN), the phosphorylated form of NR, was significantly increased upon NR supplementation (Figure 3E). Nicotinamide (NAM) and ADP-ribose (ADPR), metabolites produced by non-redox NAD^+^ dependent enzymes, were similar between NR and vehicle supplemented mice, suggesting unchanged activities of NAD^+^ consuming enzymes (Figure 3F, G). In contrast, NR supplementation significantly increased the concentration of the NAM clearance pathways products NAM oxide, N-methyl-2-pyridone-5-carboxamide (Me-2-PY), and N-methyl-4-pyridone-5-carboxamide (Me-4-PY) (Figure 3H-J). NR supplementation did not influence nicotinamide adenine dinucleotide phosphate (NADP), nicotinic acid (NA), and nicotinic acid riboside (NAR) levels (Supplemental Figure 2C-E). These data demonstrate that NR was readily utilized to improve NAD^+^ metabolism and that the subsequent elimination of NAM via methylation was enhanced in the brain (Figure 3K).

### NR alleviates behavioural deficits in a lactacystin-induced PD model

To evaluate the effect of NR on the progression of lactacystin-induced nigrostriatal neurodegeneration, we performed longitudinal behavioural tests commonly used to study the degree of unilateral neurotoxin-induced damage in mouse PD models, as indicated on Figure 3A. The elevated body swing test showed that NR significantly improved performance in the lactacystin-induced biased swing activity with the direction contralateral to the lesioned side, especially 2 weeks after lactacystin injections (Figure 4A). The cylinder test revealed that there was a significant protection against the number of contralateral rotations caused by lactacystin unilateral lesion upon NR supplementation 2 weeks after lactacystin injections (Figure 4B). When sensorimotor deficits were evaluated with the adhesive removal test, we found that the retention time to remove the adhesive from the paw at the lesioned side was superior in NR supplemented mice at 6-week time point (Figure 4C). However, the measurement of spontaneous locomotor activity using an open field test revealed a progressive decline in the total distance travelled by NR supplemented mice at the 10-week time point (Figure 4D). Altogether, our findings suggest that NR alleviates most of the behavioural deficits induced by unilateral supranigral lactacystin injection during weeks 2 to 6, but induces late onset decline in motor activity at week 10 in the open field test.

**Figure 4.**
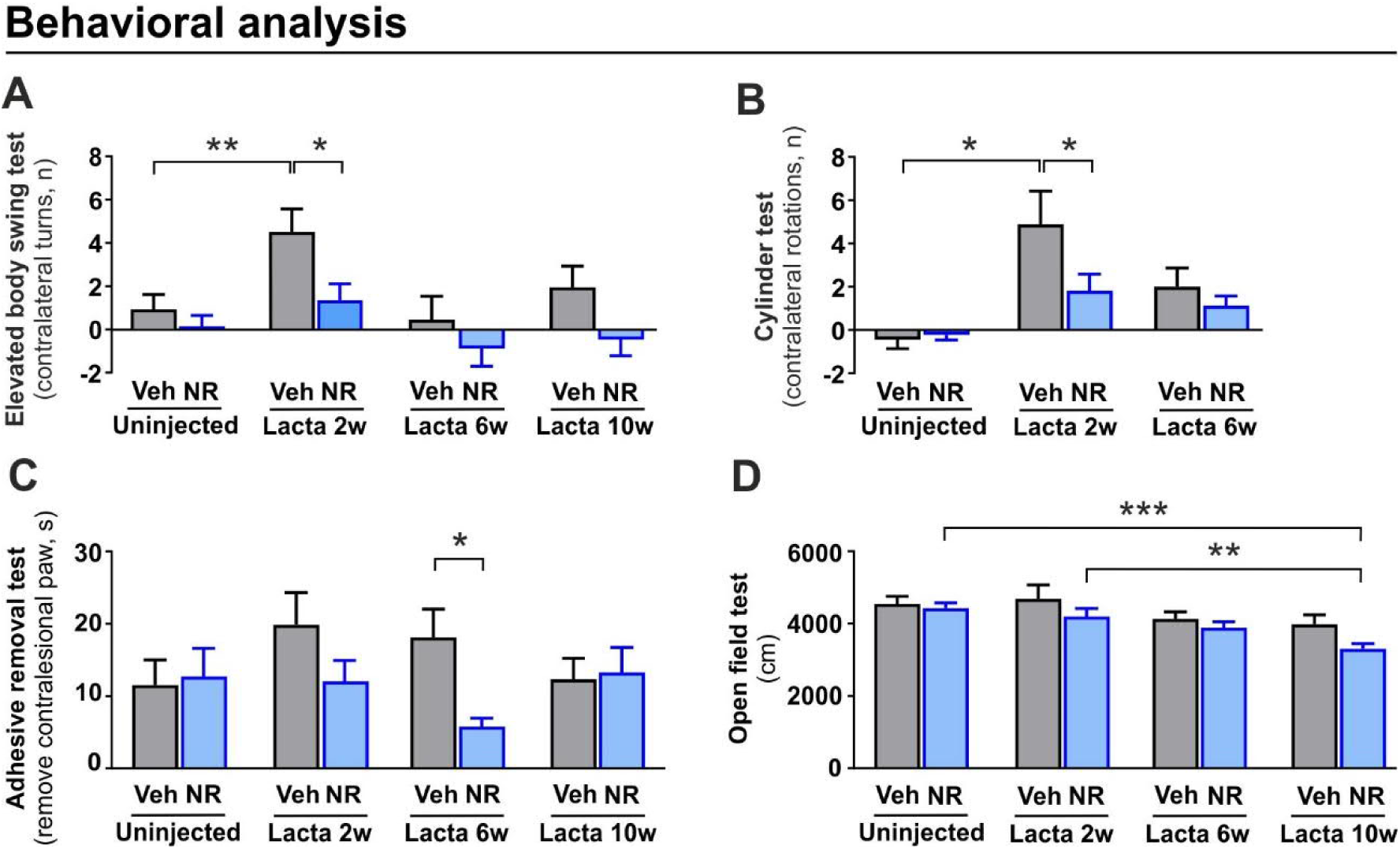
NR rescues behavioural deficits in lactacystin-induced PD model but causes a mild and gradual decline in motor activity. (**A**) Effect of NR on the number of contralateral net turns during the elevated body swing test, measured at baseline (uninjected) and 2, 6, and 10 weeks after lactacystin injection in vehicle and NR supplemented mice. (**B**) Effect of NR on the number of contralateral rotations during the cylinder test, measured at baseline (uninjected) as well as 2 and 6 weeks after lactacystin injection in vehicle and NR supplemented mice. (**C**) Time required to remove the adhesive from the contralesional paw (time-to-remove—time-to-contact) during the adhesive removal test, measured at baseline (uninjected) and 2, 6, and 10 weeks after lactacystin injection in vehicle and NR supplemented mice. (**D**) Effect of NR on the total distance travelled during the open field test, measured at baseline (uninjected) and 2, 6, and 10 weeks after lactacystin injection in vehicle and NR supplemented mice. (**A, C, D**) Veh (n = 38) and NR (n = 38) before lactacystin injection; Veh (n = 34) and NR (n = 38) after lactacystin injection. (**B**) Veh (n = 18) and NR (n = 18). Data are presented as mean ± SEM. *P < 0.05, **P ≤ 0.01, ***P ≤ 0.001. Statistical analysis was performed in GraphPad prism version 8.0.2. Comparisons between more than two groups were performed with two-way repeated measures ANOVA followed by Sidak’s multiple comparison test (**A-D**). Lacta, lactacystin; NR, nicotinamide riboside; Veh, vehicle.

### NR rescues mitochondrial dysfunction caused by nigral lactacystin lesion

To study whether NR alleviates lactacystin-induced behavioural deficits by improving mitochondrial function, we performed a second independent experiment to analyse mitochondria-related parameters 2 weeks after lactacystin injections, the time point with the clearest positive NR treatment effects on lactacystin-induced defects in the elevated body swing and cylinder tests. The study design is depicted in Figure 5A. We used high-resolution respirometry to study the effect of NR on complex I (CI), complex II (CII), and complex IV (CIV) – mediated mitochondrial respiration in unlesioned and lactacystin-lesioned nigral tissues. We did not observe significant changes in the studied parameters between vehicle and NR supplemented unlesioned side (Figure 5B, C, D). Lactacystin alone gave the expected significant drop in mitochondrial oxygen flux driven by all complexes examined in lesioned nigral tissue as compared to vehicle supplemented unlesioned side (Figure 5B, C, D). NR supplementation partially suppressed this lactacystin-induced mitochondrial bioenergetic deficit in lesioned substantia nigra (Figure 5B, C, D).

**Figure 5.**
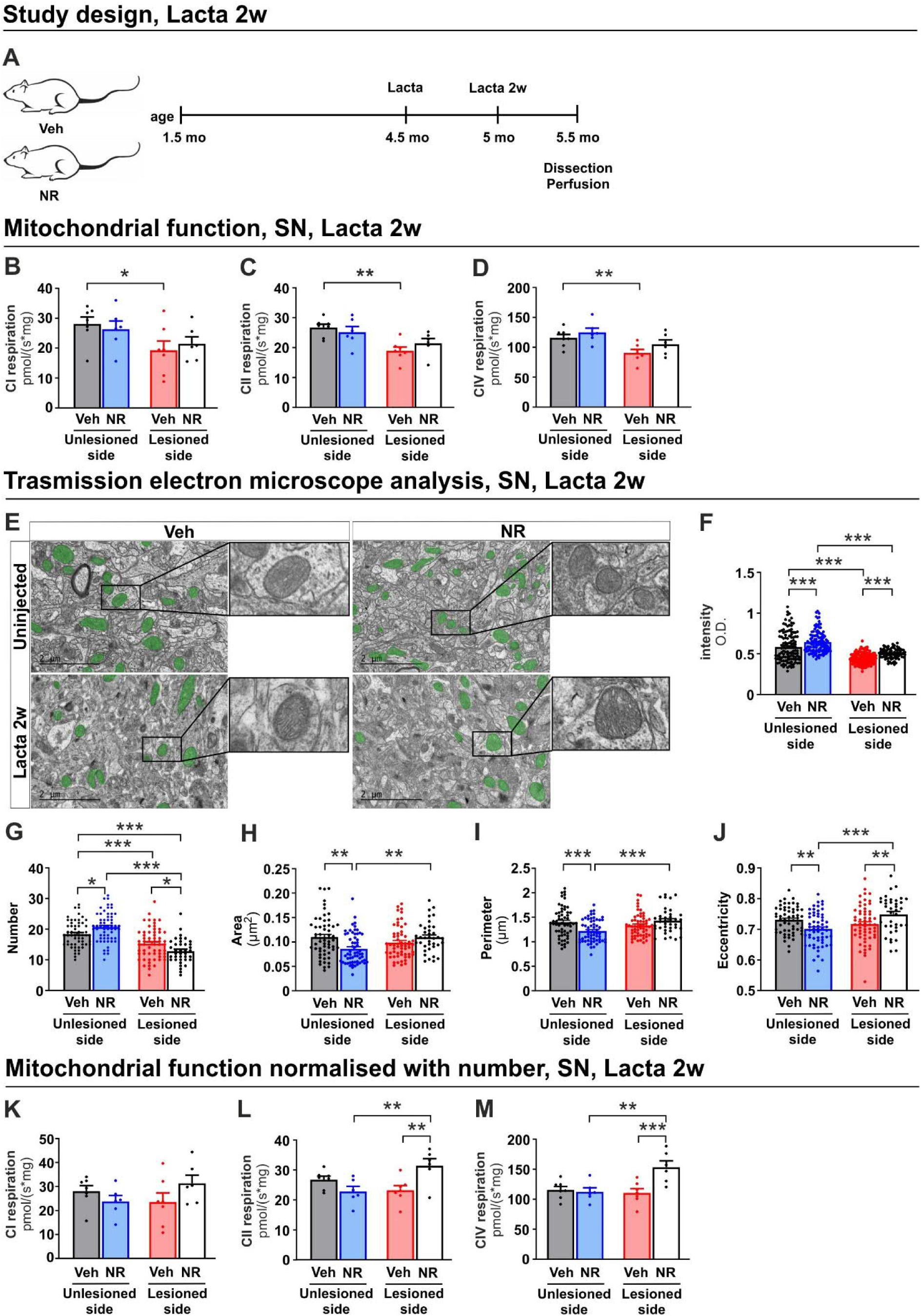
NR rescues lactacystin-induced mitochondrial dysfunction. (**A**) Experimental design for the study in which mice receiving the vehicle diet (Veh) and NR- enriched diet (NR) were analyzed 2 weeks after nigral lactacystin lesion. (**B**) Complex I [CI], (**C**) complex II [CII], and (**D**) Complex IV [CIV] mediated oxygen flux in substantia nigra tissues analyzed at the 2-week time point. Veh, unlesioned side (n = 7); NR, unlesioned side (n = 6); Veh, lesioned side (n = 7); NR, lesioned side (n = 6). (**E**) Representative transmission electron microscopy image and higher magnification inserts of the substantia nigra of vehicle and NR supplemented unlesioned and lactacystin-lesioned mice. Mitochondria are highlighted with green colour. Scale bar, 2 μm. Corresponding quantification of (**F**) mitochondria intensity, mean mitochondrial cross-section (**G**) number, (**H**) area, (**I**) perimeter, and (**J**) length/width ratio expressed as eccentricity in vehicle and NR supplemented unlesioned and lactacystin-lesioned mice. n = 3/group. Twenty non-overlapping images/sample were taken from each animal. (**K-M**) Oxygen flux data from (**B-D**) were normalized to the number of mitochondria from (**G**). Data are presented as mean ± SEM. *P < 0.05, **P ≤ 0.01, ***P ≤ 0.001. Statistical analysis was performed using GraphPad Prism version 8.0.2. Overall differences between conditions were assessed with one-way ANOVA followed by an uncorrected version of Fisher’s LSD test. CI, complex I; CII, complex II; CIV, complex IV; Lacta, lactacystin; NR, nicotinamide riboside; SN, substantia nigra; Veh, vehicle.

Given that mitochondrial function is intimately related to mitochondrial number and morphology, we carried out transmission electron microscopy (TEM) analysis of mitochondria in the substantia nigra (Figure 5E). We found that NR increased mitochondrial intensity, or darkness, an indicator of their metabolic state (33), in unlesioned side (Figure 5F). Lactacystin-lesioned nigral tissue showed significantly decreased mitochondrial intensity, which was partially rescued by NR supplementation (Figure 5F). The analysis of morphometric parameters of mitochondria revealed that NR supplemented unlesioned side had an increased abundance of mitochondria and exhibited a decrease in mitochondrial area, perimeter, and eccentricity (*i.e.,* circularity, an indicator of the mitochondrial shape defined by the length-to-width ratio) as compared to vehicle supplemented unlesioned side (Figure 5G-J). Lactacystin injection significantly reduced the number of mitochondria without a concomitant change in the mitochondrial size or shape in comparison to vehicle supplemented unlesioned side (Figure 5G, H, I). Interestingly, and contrary to the results from the unlesioned side, NR supplementation promoted a decrease in mitochondrial abundance and an increase in mitochondrial eccentricity within the lactacystin-lesioned nigral tissue (Figure 5G-J), the latter of which suggests the restoration of the normal mitochondrial circular shape in the substantia nigra.

Seeing as these mitochondrial morphology data indicate overall that lactacystin and NR drive alterations in mitochondrial dynamics, especially with respect to the amount of mitochondria, we normalized the respirometry results with the mitochondrial number obtained from TEM analysis. After this correction, it became apparent that NR significantly increased mitochondrial CII- and CIV-mediated oxygen flux per mitochondrion after lactacystin injection when compared to vehicle-supplemented mice (Figure 5K-M). The disappearance of a lactacystin-induced mitochondrial bioenergetic deficit in lesioned substantia nigra upon normalization suggests that this deficit is linked to a decline in mitochondrial number (Figure 5K-M). Taken together, these findings indicate that NR increases the functional capacity of mitochondria in lactacystin-lesioned nigral tissue at 2 weeks after the lesion.

### Long-term NR treatment decreases nigral DA level and DA related gene expression in lactacystin induced mouse model of PD

Next, we asked how NR affects DA levels in the substantia nigra and striatum of lactacystin-lesioned mice. DA levels were measured 12 weeks after nigral lactacystin lesion using high performance liquid chromatography (HPLC). In the substantia nigra, lactacystin caused a significant drop in DA tissue level when compared to vehicle supplemented unlesioned side (Figure 6A). Surprisingly, NR treatment resulted in a decrease in DA tissue levels in both the unlesioned and lactacystin-lesioned sides of the midbrain substantia nigra (Figure 6A). In the striatum, DA levels were decreased by about 60-70% in the lactacystin-injected hemisphere in both vehicle and NR supplemented mice, indicating a loss of DA function caused by nigral lactacystin lesion (Figure 6B) (28, 29). In conclusion, NR induced the loss of DA levels in both lesioned and unlesioned sides of the substantia nigra. In the striatum, NR did not prevent or exacerbate the effect of lactacystin on DA levels in the lactacystin lesioned side.

**Figure 6.**
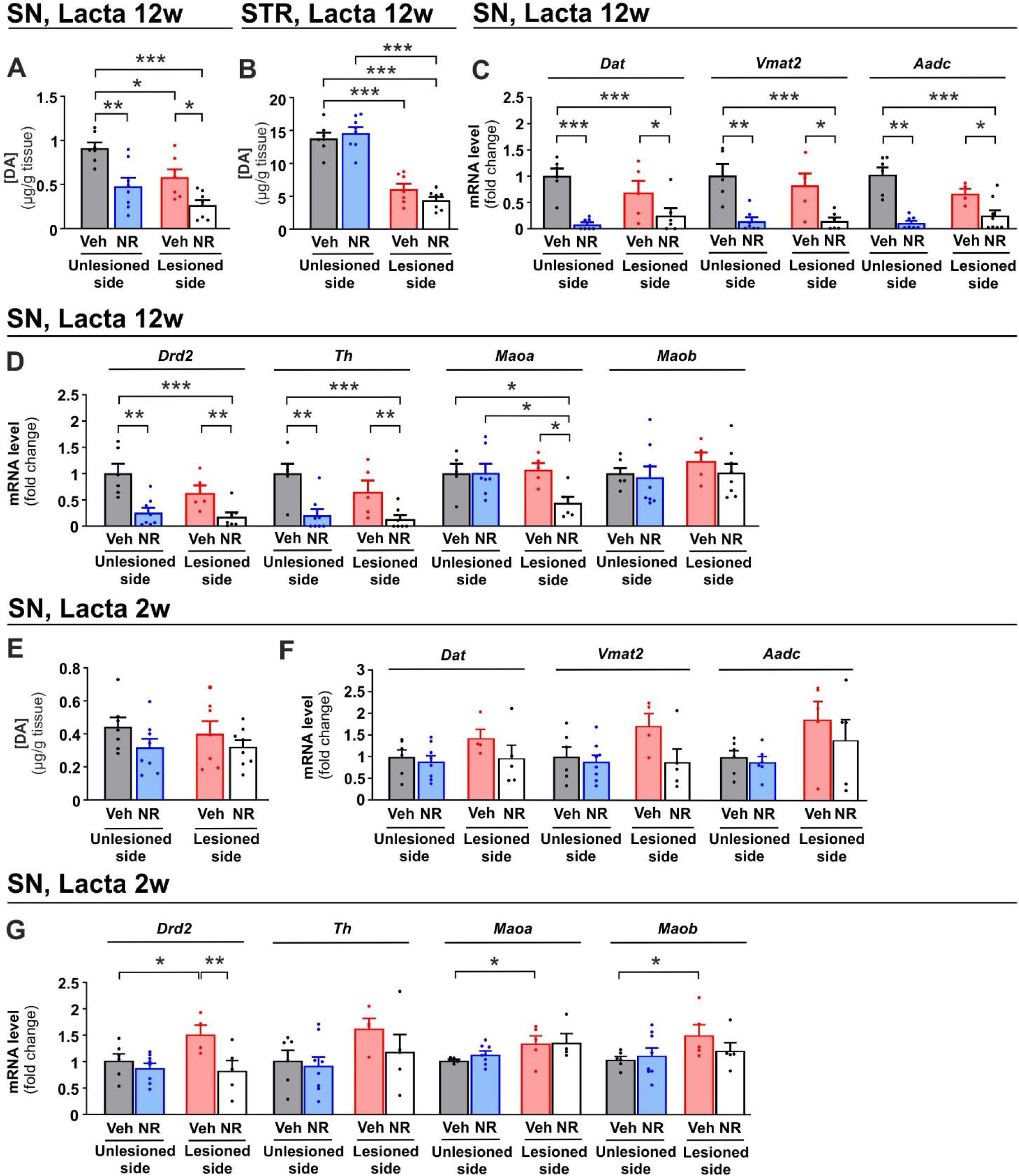
Long-term NR treatment decreases DA levels and expression of DA related genes in the substantia nigra upon lactacystin injection. Total tissue level of DA in **(A)** substantia nigra and **(B)** striatum 12 weeks after lactacystin injection. Veh, unlesioned side (n = 7); NR, unlesioned side (n = 8); Veh, lesioned side (n = 7); NR, lesioned side (n = 8). Gene expression of (C) *Dat*, *Vmat2,* and *Aadc* and (D) *Drd2*, *Th*, *Maoa*, and *Maob* mRNA in the substantia nigra of vehicle and NR supplemented unlesioned and lactacystin-lesioned mice 12 weeks after lactacystin injection. Veh, unlesioned side (n = 6); NR, unlesioned side (n = 8); Veh, lesioned side (n = 6); NR, lesioned side (n = 8). (**E**) DA tissue level in the substantia nigra of vehicle and NR supplemented mice 2 weeks after lactacystin injection. Veh, unlesioned side (n = 7); NR, unlesioned side (n = 8); Veh, lesioned side (n = 7); NR, lesioned side (n = 8). Gene expression of (**F**) *Dat*, *Vmat2,* and *Aadc* and (**G**) *Drd2*, *Th*, *Maoa*, and *Maob* in the substantia nigra of vehicle and NR supplemented mice 2 weeks after lactacystin injection. Veh, unlesioned side (n = 6); NR, unlesioned side (n = 8); Veh, lesioned side (n = 5); NR, lesioned side (n = 5). Data are presented as mean ± SEM. *P < 0.05, **P ≤ 0.01, ***P ≤ 0.001. Statistical analysis was performed using GraphPad Prism version 8.0.2. Overall differences between conditions were assessed with one-way ANOVA followed by an uncorrected version of Fisher’s LSD test. Aadc, Aromatic L-amino acid decarboxylase; DA, dopamine; Dat, dopamine active transporter; Drd2, dopamine receptor 2; Lacta, lactacystin; Maoa monoamine oxidase A; Maob, monoamine oxidase; NR, nicotinamide riboside; SN, substantia nigra; STR, striatum; Th, tyrosine hydroxylase; Veh, vehicle; Vmat2, vesicular monoamine transporter 2.

These unexpected results prompted us to analyze DA metabolism in greater detail. We first analysed the level of dopamine active transporter (*Dat*), vesicular monoamine transporter 2 (*Vmat2*), and L-DOPA converting enzyme aromatic L-amino acid decarboxylase (*Aadc*), whose activity can be assessed in clinic using positron emission tomography (PET) or single-photon emission computed tomography (SPECT) imaging to monitor PD progression (34–37). We found that the expression level of all three genes encoding for DAT, VMAT2, and AADC were downregulated in the substantia nigra in unlesioned and lactacystin-lesioned NR supplemented mice (Figure 6C). Next, we analyzed the expression of other key genes involved in DA metabolism, including DA receptor 2 (*Drd2*), tyrosine hydroxylase (*Th*), and monoamine oxidases A (*Maoa*) and B (*Maob*) in nigral tissues. We observed a significant decrease in the expression level of *Drd2* and *Th* in the substantia nigra of NR supplemented mice in both the unlesioned and lactacystin-lesioned sides (Figure 6D). In addition, *Maoa* was found to be significantly lower in lactacystin injected NR supplemented mice, whereas *Maob* expression level was unchanged (Figure 6D). Analysis of the same genes in the ventral tegmental area (VTA), the dopaminergic nucleus which connects the left (unlesioned) and the right (lesioned) substantia nigra, revealed no difference between control and NR supplemented mice (Supplemental Figure 3A), suggesting that the downregulation of DA genes by NR is specific to the substantia nigra. Analysis of genes involved in other cellular functions, including unfolded protein response of the endoplasmic reticulum (UPR^er^) as well as mitochondrial stress and quality control in the substantia nigra, revealed no changes between vehicle and NR supplemented mice (Supplemental Figure 3B, C). Collectively, these results suggest that NR supplementation results in specific downregulation of DA and DA metabolic genes in both the lesioned and unlesioned sides of the substantia nigra upon unilateral lactacystin-induced proteasome inhibition.

Next, we studied the effect of NR on tissue DA levels and on the expression of key genes involved in DA metabolism 2 weeks after lactacystin injection, the time point at which NR markedly rescued lactacystin-induced behavioural deficits (Figure 5A). We observed no significant change in DA levels in the substantia nigra of both unlesioned and lactacystin-lesioned sides at this time point (Figure 6E). The levels of *Dat*, *Vmat2*, and *Aadc* also did not differ between vehicle and NR supplemented groups 2 weeks after lactacystin injection (Figure 6F). However, lactacystin significantly increased *Drd2*, *Maoa*, and *Maob* mRNA levels in lesioned nigral tissue. Such an upregulation of *Drd2*, *Maoa*, and *Maob* was not observed in the NR supplemented group (Figure 6G). In contrast to the lactacystin lesioned animals, NR had no effect on tissue DA levels or on the expression of DA metabolism related genes in the substantia nigra of mice that did not undergo lactacystin injection (Supplemental Figure 4A-C).

Collectively, these data suggest that despite the improved mitochondrial function and behavioural phenotypes, NR does not rescue tissue DA levels in the striatum and the substantia nigra. Instead, NR induces bilateral downregulation of tissue DA and DA metabolism related genes 12 weeks after unilateral proteasome inhibitor lactacystin injection in the substantia nigra.

## Discussion

The overall success of NR in enhancing mitochondrial function, muscle performance, and metabolic health in several disease models (38) has led to rapid implementation of NR into clinical trials of various diseases, including PD (NCT03816020, NCT03568968, NCT04044131). However, few supporting studies are performed in preclinical PD models. Hence, our understanding of how NR influences PD-relevant endpoints such as mitochondrial function, proteostasis, and DA cell function is limited. Our study addresses this important knowledge gap and reveals that NR provides functional benefit by restoring the mitochondrial function and proteostasis in both α-syn overexpressing *C. elegans* models and - at first sight - in a lactacystin mouse model of PD. However, we found that long-term NR application causes an adverse effect in lactacystin induced 26S/20S proteasome inhibition model of PD in mice: a decrease in DA level and downregulation of essential DA metabolic genes in both the lesioned and unlesioned side of the substantia nigra. Our results call for caution in clinical trials, invite for further experiments in various PD models and suggest specific endpoints to examine in the ongoing clinical trials with NR.

Our studies using *C. elegans* PD models with muscle- or DA neuron-overexpressing α-syn demonstrate that these animals recapitulate typical features of α-syn neurotoxicity, including DA neuron and motor coordination loss as well as mitochondrial dysfunction. In agreement with previous publications showing disturbed mitochondrial proteostasis in PD (10, 11), we detected a significantly decreased expression of multiple UPR^mt^ -associated mitochondrial protease genes in the muscle α-syn *C. elegans* model, possibly implying the exhaustion of UPR^mt^ signalling. Collectively, our data reveal that the loss of UPR^mt^ signalling and mitochondrial proteostasis likely contribute to the molecular mechanisms underlying the α-syn mediated mitochondrial dysfunction. Conversely, increasing mitochondrial function and UPR^mt^ has emerged as a potential approach to modulate neurodegenerative diseases such as Alzheimer’s disease (20). The supplementation with NR likely restored the α-syn overexpression driven mitochondrial dysfunction, proteostasis failure, and the motility defect via *atfs*-*1*-mediated UPR^mt^ activation. Altogether, the *C. elegans* data encouraged further studies on long-term NR supplementation in mice.

To that end, we used the proteasome inhibitor lactacystin supranigral delivery in mice to model PD (28, 29). Lactacystin-induced inhibition of 26S/20S proteasomal function causes an upregulation of α-syn expression (29), mitochondrial dysfunction, and cell death *in vitro* (39) and *in vivo* (40). Similarly, α-syn overexpression leads to disturbances in the proteasome activity in rodents (41) and compromises the 26S/20S proteasome function in the substantia nigra in PD patients (4, 5). In line with our results from *C. elegans,* NR improved mitochondrial function in the substantia nigra in mice. This finding is in agreement with various previous mouse studies showing NR’s capacity to boost mitochondrial function (42, 43). Moreover, NR rescued the behavioural deficits caused by nigral lactacystin lesion. Interestingly, NR supplementation led to the peculiar result of a divergence between behavioural and mitochondrial phenotypes, which were almost universally improved, and DA metabolism, which was impaired. We observed a time-dependent reduction in tissue DA levels and the downregulation of the key DA metabolic genes in both the lesioned and unlesioned substantia nigra in lactacystin injected mice upon NR. These findings were paralleled with a decrease in spontaneous locomotor activity in NR supplemented mice. As NR did not influence striatal or nigral DA levels at the 2-week time point, when the NR-induced rescue of lactacystin induced behavioural deficits was the most prominent, the observed beneficial effect of NR is likely due to the improvement in mitochondrial function and not because of enhanced DA production. Importantly, our study revealed that NR alone, without unilateral supranigral lactacystin delivery, does not affect nigral DA levels and DA metabolism and NR-induced DA metabolism deterioration occurs after the 2-week and before the 12-week time point in lactacystin induced PD model. Collectively, NR failed to protect or restore loss of nigrostriatal DA and DA metabolism, one of the most important endpoints in PD preclinical and clinical trials (44), upon proteosome inhibition.

A potential mechanism via which NR could influence DA metabolism is the effect of supplementation on cellular methylation capacity (45). When taken as an oral supplement, NR is converted to the entire NAD metabolome and subsequently degraded to NAM (32), which is eliminated via methylation to MeNAM, Me-2-PY, and Me-4-PY, excreted in the urine. We found that methylated NAM waste products were significantly elevated in the cerebellum of NR supplemented mice. Conceivably, chronic activation of the NAM methylation process catalysed by NNMT could result in depletion of the methyl donor *S*-adenosylmethionine (SAM) utilized in this reaction. The deficiency of SAM could then limit synthesis of DA driven by tyrosine hydroxylase (TH) (46) and influence the transcription of DA metabolism related genes by reducing DNA methylation (47) in lactacystin-injected mice. Given that DNA hypomethylation has been reported in post-mortem brain samples of PD patients (48), NR supplementation in PD models or patients might benefit from the support of a methyl donor such as folic acid or trimethylglycine. In principle, NR could have suppressed expression of DA metabolism related genes by increasing the levels of NAM (32) as NAM has been reported to exacerbate DA neuron degeneration in a lactacystin rat model for PD (49) and decreased the number of TH-positive DA neurons in neonatal mice (50). However, brain NAM content remained unaltered upon NR supplementation, excluding the possibility that NAM mediated neurotoxicity drives the downregulation of DA metabolism in the substantia nigra in our study. So far, the longest clinical trial conducted with NR addressing its effect on mitochondrial function and metabolic health have been limited to 12 weeks (51–53). Our results suggest that long term NR application may exert negative effect on dopamine metabolism upon proteasome inhibition. Whether 26S/20S inhibition induced by lactacystin in mouse substantia nigra resembles 26S/20S inhibition observed in post-mortem substantia nigra in PD patients (4, 5) remains an important topic of future research. Nevertheless, current results imply that long term NR application in PD patients, a sub-group of PD patients or any individual with genetic or environmental reduction in 26S/20S function, may have adverse effects. For ongoing PD clinical trials, our results argue for collecting additional safety data, including, SPECT and/or PET imaging of DAT, VMAT, or AADC. Our observation that NR supplemented PD mice display a gradual reduction in open field motor activity also suggest that the monitoring of individual longitudinal motor scores during the trial may be especially relevant. Collectively, deeper and longer-term studies in a variety of PD models to better evaluate the potential long-term effects of NR are urgently needed.

## Material and Methods

### *C. elegans* growth and maintenance

*C. elegans* strains were cultured at 20°C on nematode growth medium (NGM) agar plates, seeded with E. coli strain OP50 as a food source. Strains used in this study were the wild-type Bristol N2, DDP1 (*uonEx1*[*unc-54*p*::α-synuclein::CFP* + *unc-54p::α-synuclein::YFP*(Venus)], which overexpresses wild-type α-synuclein in body wall muscle cells (21) and is referred to here as muscle α-syn, DDP2 uonEx2 [*unc-54p*::CFP::YFP(Venus)] (54), the control strain for DDP1, and BY273: Is[P*dat-1::*GFP; P*dat-1::*α-syn] (18), which overexpresses wild-type α-synuclein in dopamine (DA) neurons and is referred to here as the DA α-syn strain. The adult muscle α-syn animals exhibit a significantly lower pharyngeal pumping and locomotion rate than wild-type animals (21). In turn, the DA α-syn strain exhibits age-associated dopaminergic neuron degeneration (22). The strains were provided by the Caenorhabditis Genetics Center (CGC, University of Minnesota). In order to avoid any falsifying effect on cellular metabolism, mitochondrial stress (20), and overall life expectancy (21), *C. elegans* experiments were performed without an addition of 5-fluoro-2’-deoxyuridine, a DNA synthesis blocking drug commonly used to sterilize *C. elegans.* The bleach-treated, age-synchronised population was maintained in NGM plates by manually transferring the animals to fresh plates every second day until the desired age was attained.

### Treatment of *C. elegans*

NR chloride (ChromaDex Inc, USA) dissolved in water and water serving as a vehicle were added on top of the plate and swirled over to cover the entire plate. NR was used at a final concentration of 1 mM. The plates were left to air dry for 2 hours before transferring *C. elegans* onto them. In all the experiments, *C. elegans* was exposed to NR from the L1 larval stage and continuously throughout the experiment, unless stated otherwise. To ensure constant exposure to the compound, plates were changed every second or third day.

### Oxygen consumption assay

Oxygen consumption rate was measured using the Seahorse XF96 respirometer (Agilent Bioscience) as described in (55). For each experiment, at least 10 to 15 wells containing 200 µL of M9 buffer were used per group, with 5 to 8 *C. elegans* being transferred manually into each well. After the measurement of five cycles of basal respiration, carbonyl cyanide-4-(trifluoromethoxy)phenylhydrazone (FCCP) at a final concentration of 15 µM was injected and another five cycles of maximal respiration were measured. Basal and maximum respiration rates were normalized to the number of animals in each individual well. The average values of five cycles of basal and maximal respiration were calculated. Each experiment was repeated at least twice.

### Analysis of mitochondrial DNA amount

Quantitative real time PCR (qRT-PCR) was used to determine the mitochondrial DNA (mtDNA) amount in the muscle α-syn *C. elegans* strain. DNA was extracted with a phenol–chloroform extraction, followed by ethanol and sodium acetate precipitation (56). Two different mitochondrial genome regions (*nd-1* and *mtce.26)* and one nuclear region (*act-3)* were amplified using the gene region-specific primers. The mtDNA amount was presented as a relative level of mtDNA genome per nuclear genome. The qRT-PCR reaction was performed similarly, as mentioned in the gene expression analysis found in the method section ‘gene expression analysis in *C. elegan*s,’ except the template amount used in this instance was 400 pg of DNA per well. Data analysis was performed using the standard curve method, which utilizesthe qBASE+ 3.2 software (Biogazelle). Each experiment was repeated twice and was performed on at least three independent biological samples. Primer sequences are provided in the Supplemental Table 1.

### Pharyngeal pumping

The recently developed microfluidic-based ScreenChip system (NemaMatrix, USA) was used to monitor the pharyngeal pumping rate in *C. elegans* (57). For this assay, animals were grown in an age synchronized manner. The *C. elegans* were washed twice with an M9 buffer before they were incubated with 10 mM of serotonin at room temperature for 20 minutes. Simultaneously, the NemaMatrix fluidics system was prepared according to the manufacturer’s instructions. Then, the animals were loaded into a 1.5 ml tube and sucked into the ScreenChip through a fine tubing and vacuum pump system. To achieve correctly positioned *C. elegans*, the pumping frequency was measured by utilizing the electropharyngeogram recording of NemaMatrix ScreenChips 40 and 60 for young and old *C. elegans,* respectively. Each recording was 1 to 2 minutes long, with 15 to 20 animals per strain per group.

### Motility assay

*C. elegans* movement (*i.e.* motility analysis) was performed as described in (58). Briefly, the NGM plates containing the nematodes (with 15 to 20 animals per plate) were gently tapped, and the movement was recorded under an inverted stereomicroscope. The motility was then tracked by an automated tracker using MATLAB software. The result was quantified as the total distance normalised to the size of the animals. The experiments were repeated at least twice.

### Measurement of DA neuron survival and neurite length

The degeneration of DA neurons in the DA α-syn *C. elegans* strain was monitored by quantifying green fluorescent protein (GFP) signal intensity under the dopamine transporter (*dat-1*) promoter. For NR supplementation, DA α-syn animals were exposed to 1 mM of NR at the L4 larval stage. Approximately 15 to 20 adult animals were immobilized with 2 mM of levamisole submerged in M9 buffer on the agarose pad per each time point analyzed. The DA neurons were imaged under a Zeiss LSM780 confocal microscope with the Zeiss Plan-Apochromata 63x 1.4 numerical aperture objective. GFP fluorescent intensities, as well as background, were quantified using the Fiji ImageJ software. The threshold was selected from the brightest image and this same threshold was applied to all images from the same experiment. The average intensity of DA neurons consisting of cephalic ‘CEP’ and posterior deirid ‘PDE’ neurons was analyzed. Neurite length was also measured using ImageJ. The plugin of ImageJ for the measurement of simple neurite anatomy was used to quantify the length of each neuron in µm.

### Eight-OH-dG immunohistochemistry

Young and old *C. elegans* were freeze-cracked on dry ice blocks. Immunostaining was performed as described previously (59, 60). *C. elegans* on the slides were briefly fixed in acetone and methanol at a 1:1 ratio for 10 minutes at −20°C and were subsequently washed in PBS-T (1 x PBS, 0.1% Tween-20) for 5 minutes, followed by 30 minutes of incubation with an image-IT FX signal enhancer (Invitrogen) and 30 minutes of blocking in PBS-TB (1 x PBS, 0.1% Tween-20, 0.5% BSA). The slides were incubated with a primary antibody overnight at 4°C and washed three times in PBS-T for 10 minutes each, followed by a period of incubation with the secondary antibody at room temperature for 2 hours. The 8-OH-dG primary antibody (4354-MC-050 Trevigen) was used with a 1:200 dilution. The secondary antibody used was the Alexa Fluor 555-conjugated anti-mouse (1:1500). The samples were mounted with the Prolong Gold Mountant. The fluorescent stain 4′,6-diamidino-2-phenylindole (DAPI) (Invitrogen, P36931) and GFP were used to detect nuclei and α-syn accumulations, respectively. The samples were imaged with a Zeiss LSM780 confocal microscope equipped with the Zeiss Plan-Apochromat 63x 1.4 numerical aperture objective.

The quantification for 8-OH-dG in DA α-syn strain was performed by counting the number of 8-OH-dG positive foci in the head region of the animals. Given that the number of 8-OH-dG positive foci was remarkably high in muscle α-syn strain, beyond the countable range, 8-OH-dG staining fluorescence intensity was measured with procedures similar to those described above in the ‘DA neuronal survival assay’.

### *C. elegans* RNA interference

RNA interference (RNAi) was performed using the feeding protocol described previously (61). Briefly, the HT115 bacterial strain carrying the empty pL4440 expression vector was used as a control in all experiments. Double-stranded RNA expression was induced by the addition of 0.4 mM of isopropyl-β-D-thiogalactopyranoside (I6758, Sigma, St. Louis, MO, USA) during peak culture growth, and its concentration was increased further to 0.8 mM just before the seeding of the feeding plates. Age-synchronized L4 larvae were placed on control and RNAi-seeded plates, targeting *atfs-1* (ZC376.7, Source BioScience, J. Ahringer library).

### Mice and their maintenance

This study was performed using C57BL/6 male mice. Mice were maintained at temperature-controlled conditions of 20 to 22°C under a 12hours/12hours light/dark cycle at a relative humidity of 50-60%. Cages and bedding material (aspen chips, Tapvei Oy, Finland) were changed every week, and wooden tube and aspen shavings were provided as enrichment. Mice received food and water *ad libidum* and body weight was measured every second week. For food intake measurements, pellets were weighed before and after food supplementation and the difference in weight divided by the number of mice in the cage was taken as a measurement of overall intake. Two independent measurements were done.

All experiments were conducted following the 3R principles of the EU directive 2010/63/EU, which govern the care and use of experimental animals, and were approved by the County Administrative Board of Southern Finland (licence numbers ESAVI-2010-09011/Ym-23 and ESAVI/11198/04.10.07/2014). The protocols were authorized by the national Animal Experiment Board of Finland.

### Study design

Mice were randomly divided into two groups and fed with a chow diet containing vehicle (water) or NR (400 mg/kg/day) (42). The supplementation with NR was started when mice were 6- to 7-weeks-old. Pellets were prepared by mixing powdered chow diet (2016S, Harlan Laboratories) with water or NR dissolved in water and were dried under a laminar flow hood at room temperature for 48 to 72 hours and stored at −20°C, as described previously (20, 42, 62, 63). Fresh vehicle and NR-containing food pellets were provided to mice every week. To evaluate the effect of NR on the progression of nigrostriatal neurodegeneration, mice were preconditioned with vehicle or NR for 3 months and the behavioural phenotype was assessed 2 weeks before and 2, 6, and 10 weeks after unilateral lesion in the substantia nigra. We performed two independent experiments. In the first experiment (Figure 3A), mice were dissected 12 weeks after nigral lesion; in the second experiment (Figure 5A), mice were dissected 2 weeks after nigral lesion. In the second experiment, a group of mice that received vehicle and NR diets but did not undergo nigral injection was also included (Supplemental Figure 4A). For tissue collection, mice were anaesthetized with CO2 and euthanized by cervical dislocation followed by decapitation. The brain was quickly removed from the skull, immersed in ice-cold PBS, and placed in a brain block (Stoelting, Wood Dale, IL). Brain regions of interest were collected as described in (28) using a puncher (inner diameter 2mm). They were then snap frozen and stored at −80°C until they were ready to be processed. To induce neurodegeneration, we used the 26S/20S proteasome inhibitor lactacystin model of PD, where lactacystin is unilaterally injected just above the nigrostriatal DA neurons in the substantia nigra. We chose the lactacystin model over dopamine neuron-specific toxin-induced PD models such as 1-methyl-4-phenyl-1,2,3,6-tetrahydropyridine (MPTP) and 6-hydroxydopamine (6-OHDA) because specifically killing off DA neurons with the latter two toxins does not reflect general proteostasis failure in PD, while a reduction in proteasome function and the loss of 26S/20S proteasome subunits in post-mortem analyses in the substantia nigra of PD patients is well characterized (4, 5) and reviewed in (26). In cell-lines and primary dopamine neurons, lactacystin leads to the generation of α-syn oligomers and the formation of cytoplasmic inclusion bodies containing ubiquitin and α-syn (26). Further supporting our choice, our experiences, as well as results from other laboratories, show that supranigral lactacystin delivery in the used dose range results in a reproducible and consistent lesion with an approximately 40% loss of DA cells in the substantia nigra pars compacta and concurrent striatal DA depletion (28–31). This contrasts with the MPTP- and 6-OHDA-induced lesions, which often give variable results among different laboratories. Similarly, literature with α-syn-induced DA neuron degeneration is often controversial and difficult to reproduce in mice (64).

### Lactacystin injection

For lactacystin injections, mice were anesthetized with isoflurane and placed into the stereotaxic frame (Stoelting). After disinfecting the skin, 0.1 ml of lidocaine with adrenaline (10 mg/ml, Orion Pharma) were injected under the scalp. A small cut was made along the top of the head to expose the skull and burr holes were made with a high-speed drill. A 10 μl Hamilton syringe with a 26G steel needle attached was used to inject lactacystin (A.G. Scientific/Nordic Biosite) into the right brain hemisphere, just above the substantia nigra (AP −3.3; ML −1.2 and DV −4.6). The volume for each injection was 3 μl, with a flow rate of 0.5 μl/minute, and the needle was left to sit for 5 minutes after the injection. The total amount of lactacystin injected was 3 μg. After stitching the wound, carprofen (5 mg/kg s.c., Pfizer) was administered to hinder post-operative pain. Mice were returned to their home cage upon awakening. For the experiments, the non-lesioned hemisphere (left side) was used as the control. After lactacystin injection, 4 mice from the control group died.

### Elevated body swing test

Asymmetrical motor behaviour was assessed with the elevated body swing test (65). The mouse was lifted 20 times, approximately 5 cm from the base of its tail. A swing was recorded whenever the animal moved his head with a deviation of more than 45° in each direction. The animal was returned to his resting position with all four paws on the floor before being lifted up again.

### Cylinder test

Motor asymmetry was assessed with the cylinder test (66). Freely moving mice were recorded for 5 minutes in a plexiglass cylinder (diameter 20 cm) and an observer who was blind to the identity of the samples counted the number of ipsi- and contra-lateral rotations (with an amplitude of 360 °).

### Adhesive removal test

Sensorimotor deficits were assessed using the adhesive removal test (67). Mice were first acclimated to the testing box for 60 seconds. After this period, the mouse was removed from the testing box, and two adhesive round tapes were subsequently applied with equal pressure to the hairless part of each forepaw. The mouse was placed back in the testing box afterwards. The time to touch and remove the adhesive from each paw was recorded over a period of 2 minutes. The adhesive place order (right or left) was alternated and randomized within each group to reduce bias.

### Open field test

General spontaneous locomotor activity was measured using the open field test (68). The apparatus consists of an open field arena (30 × 30 cm, Med Associates, St. Albans, VT) equipped with infrared light sensors detecting horizontal and vertical activity. During the experiment, the mice were placed on the corner of the arena and spontaneous activity was recorded over a period of 30 minutes at a light intensity of 150lx. The peripheral zone was defined as a 6 cm wide corridor along the wall. Data was recorded and analyzed with Activity Monitor v5.1 software (Med Associates, St. Albans, VT).

### Measurement of mitochondrial function in tissue samples

Fresh substantia nigra tissues were rinsed with ice-cold PBS and homogenized with glass-teflon potter, 4 mg of tissue per ml of MiR05 buffer (0.5 mM EGTA, 3 mM MgCl2, 60 mM lactobionic acid, 20 mM taurine, 10 mM KH2PO4, 20 mM HEPES, 110 mM D-sucrose, 1 g/l BSA). Two mg of tissue homogenate were measured in a pre-calibrated chamber of an Oxygraph-2k (Oroboros Instruments, Innsbruck, Austria) at 37°C. Complex I-mediated oxygen flux was induced by the administration of substrates malate (5 mM), pyruvate (5 mM), and glutamate (5 mM), along with ADP (2 mM), and was inhibited with complex I inhibitor rotenone (0.15 µM) to determine the complex I-mediated oxygen flux. This was followed by an addition of the complex II substrate succinate (34 mM) and the shutdown of electron transfer by complex III inhibitor antimycin A (2.5 µM) to measure the complex II-mediated oxygen flux. Complex IV oxygen flux was measured using *N, N, N*′, N′-tetramethyl-*p*-phenylenediamine (TMPD, 0.5 mM) as a substrate, in conjunction with the reducing agent ascorbate (2 mM). Complex IV was inhibited with sodium azide (50 mM) to account for the chemical auto-oxidation. Residual oxygen flux was measured after antimycin A administration. Oxygen flux was normalised to both the tissue mass and the mitochondrial number quantified from the TEM image of the substantia nigra.

### Transmission electron microscopy

Mice were anaesthetized with a lethal dose of pentobarbital (Mebunat Vet) and transcardially perfused with PBS, followed by decapitation and the isolation of the substantia nigra. The tissue pieces (1mm x 1mm) were postfixed, stained, and embedded as described in (69).

#### Mitochondria analysis

For mitochondria analysis, 20 non-overlapping images/sample were taken at 4000x magnification using a Jem-1400 transmission electron microscope (Jeol, Tokyo, Japan). The collected images were processed, aligned, and analyzed using a Microscopy Image Browser (70) by a person blind to the identity of the samples.

#### Mitochondria Optical Density measurement

The Optical Density (O.D.) analysis of the mitochondria was performed blinded from the same images used previously. Measurements were performed using Fiji ImageJ (ImageJ, Version 1.52). Image calibration was performed using a calibrated O.D. step tablet containing 21 steps with a density range of 0.05 to 3.05 O.D. (Kodak Calibrated Step Tablet scanned with an Epson Expression 1680 Professional scanner). Background correction in each section was performed by subtracting the average of the O.D. values taken from three empty areas from the O.D. value of each mitochondrion from the same section. Mitochondria undergoing fission, fusion, or apoptosis were excluded from the analysis.

### High performance liquid chromatography

DA levels were analyzed from the dissected brain samples, as described previously (71), using High Performance Liquid Chromatography (HPLC) with electrochemical detection. The values are presented as micrograms per gram of wet tissue weight.

### NAD^+^ level quantification assays

#### C. elegans

NAD^+^ content in the muscle α-syn *C. elegans* strain was measured using an NADmed method recently developed at the University of Helsinki. It is a conventional assay based on enzymatic reaction with colorimetric detection (https://www.helsinki.fi/fi/tutkimusryhmat/mitochondrial-medicine and www.NADmed.fi).

#### Mouse

Although the majority of the analyses were conducted in the substantia nigra, the analysis of the brain NAD^+^ metabolome was performed using the cerebellum because of the large amount of tissue required for this purpose. Prior to extraction, the brain samples were pulverized in a Bessman pulverizer prechilled with liquid nitrogen. Frozen samples were weighed into two tubes with an accuracy of 0.1 mg. Two extractions were performed to determine the complete NAD^+^ metabolome (26). In the first set, NAD^+^, NADP, NMN, NAR, NAAD, and ADPR were analysed, and stable 13C labelled analogues of the measured nucleotides and neutral nucleosides were added to equally weighted tubes (72). Four hundred µL of hot buffered ethanol (25% 10 mM HEPES, 75% ethanol) were added to each tube and subsequently vortexed for 15 seconds, with the samples then being placed on ice until the processing of the batch was completed. Next, the samples were heated for 3 minutes at 55°C on a vortex shaker, followed by centrifugation at 4°C with maximum speed for 10 minutes. Supernatant was collected and evaporated on a SpeedVac until dry. Meanwhile, standards, controls, and blanks were prepared in hot buffered ethanol with the same internal standard solution, and were then dried under vacuum alongside the samples. Samples were reconstituted at the time of analysis with 97% 10 mM ammonium acetate/ 3% acetonitrile. The second set of samples was analysed for NR, NAM, NA, MeNAM, NAM oxide, Me-2-PY, and Me-4-PY prepared in the same fashion, except that the internal standard mixture contained 18O-NR, 18O-NAM, d3, 18O-MeNAM, d4-NA, and d3-Me4py. LCMS was performed as described previously (26).

### Gene expression analyses

#### C. elegans

Approximately 1500 animals per condition were collected in M9 buffer from NGM plates in three biological replicates. Total RNA was isolated using Trizol Reagent (Thermo Fisher Scientific) according to the manufacturer’s protocol. One microgram of RNA was reverse transcribed into complementary DNA using the QuantiTect Reverse Transcription Kit (Qiagen). The template amount of 6.6 ng RNA equivalent per wellt was used and the PCR reaction was performed in triplicate using the total volume of 8 µl in 384 well plate format. Gene expression analysis of selected genes was conducted using the LightCycler 480 SYBR Green I Master reagent (Thermo Fisher Scientific) in CFX384 Touch™ Real-Time PCR Detection Systems (Bio-Rad Laboratories). To normalize the expression data, two housekeeping genes were used: putative iron-sulfur cluster assembly enzyme (Y45F10D.4) and cell division cycle-related (cdc-42). Data analysis was performed with the standard curve method using the qBASE+ 3.2 software (Biogazelle). Primer sequences are provided in the Supplemental Table 1. Each experiment was repeated twice.

#### Mouse

Total RNA was isolated from frozen tissues using Trizol Reagent (Thermo Fisher Scientific) according to the manufacturer’s protocol, and RNA quantity and quality (absorbance 260/280 nm>1.8) were assessed using NanoDrop (NanoDrop Technologies). One hundred fifty nanograms of DNase I (Thermo Fisher Scientific)-treated total RNA was reverse transcribed into complementary DNA using random hexamer primers and RevertAid Reverse Transcriptase (Thermo Fisher Scientific). The template amount of 1.875 ng RNA equivalent per well was used, and the PCR reaction was performed in triplicate using the total reaction volume of 10 µl in 384 well-plate. Gene expression analysis of selected genes was conducted using the BioRad C1000 Touch Thermal Cycler, which was upgraded to the CFX384 System (Bio-Rad Laboratories). Mouse *Gapdh,* or a combination of *Gapdh*, *Actb*, and *18S rRNA,* were used as reference genes. Results for biological repeats were discarded when the Cq value for one or more of the replicates was 40 or 0, or when the Cq difference between replicates was >1. Expressions of genes were calculated with the ΔΔ*C*t method (73) and analyzed with CFX Manager^TM^ Software (Bio-Rad). Primer sequences are provided in Supplemental Table 2.

### Statistical analysis

Comparisons between two groups were analysed with Student’s t-test followed by Welch’s correction. Comparisons between more than two groups were performed with the One-way Analysis of Variance (ANOVA) test followed by Fisher’s LSD test. Outliers were identified with Grubbs’ test and were excluded from analysis. Statistical analyses were performed with PRISM 8.0 (GraphPad software).

## Author contribution

G.T., F.A., T.S., S.P.K., S.O., K.M., A.M., and V.H. performed the experiments. M.S. and C.B. performed and analysed brain NAD^+^ metabolome. P.P. performed and analysed HPLC. G.T. and F.A. prepared the figures. G.T., F.A., E.P., and J.-O.A. wrote the manuscript. C.I.H. and H.N. co-supervised the *C. elegans* experiments and provided funding. E.P. and J.-O.A. supervised the project, planned experiments, and provided funding. All the authors have read and approved the manuscript.

## Acknowledgements

G.T. was supported by the Finnish Parkinson Foundation. T.S was supported by a grant from the Research Council of Norway (grant no 302483). K.M. was supported by the doctoral program Brain and Mind and by the Alfred Kordelin Foundation. C.I.H was supported by the Academy of Finland (grant no 297776) and the Sigrid Jusélius Foundation. J.-O.A. was supported by the Academy of Finland (grant no 297727), the Sigrid Jusélius Foundation, the Faculty of Medicine at the University of Helsinki, the Helsinki Institute of Life Science (HiLIFE) Fellow grant, the European Research Council (ERC) (grant no 724922), and the Alzheimerfonden. E.P. was supported by the Academy of Finland (grant no 286359). We thank Kimmo Haimilahti, Minna Kuusela, Noora Pöllänen, Minna Tuominen, Rita Rinnankoski-Tuikka, and Anita Wagner for their valuable help and technical assistance during the mouse experiments, and we thank Sweta Jha for the assistance with *C. elegans* experiments. The authors also thank Dr. Kira Holmström and Sonja Koopal for their experimental advice when the project was initiated. The authors also wish to thank Liliya Euro for the development of NAD analysis protocol and for technical advice. We also acknowledge the Chromadex CERP Science Team for the fruitful discussion and the Electron Microscopy Unit of the Institute of Biotechnology, University of Helsinki, for providing laboratory facilities. Finally, the authors thank Ian Mitchell for language editing. The DA α-syn *C.elegans* strain was provided by Blakely Lab (FAU Brain Institute). Other *C.elegans* strains were provided by the CGC, which is funded by the NIH Office of Research Infrastructure Programs (P40 OD01C. elegans0440).

