## Supplemental Material for "Nicotinamide riboside alleviates Parkinson’s disease symptoms but downregulates dopamine metabolism upon lactacystin-induced proteostasis failure"

### SUPPLEMENTAL FIGURE 1

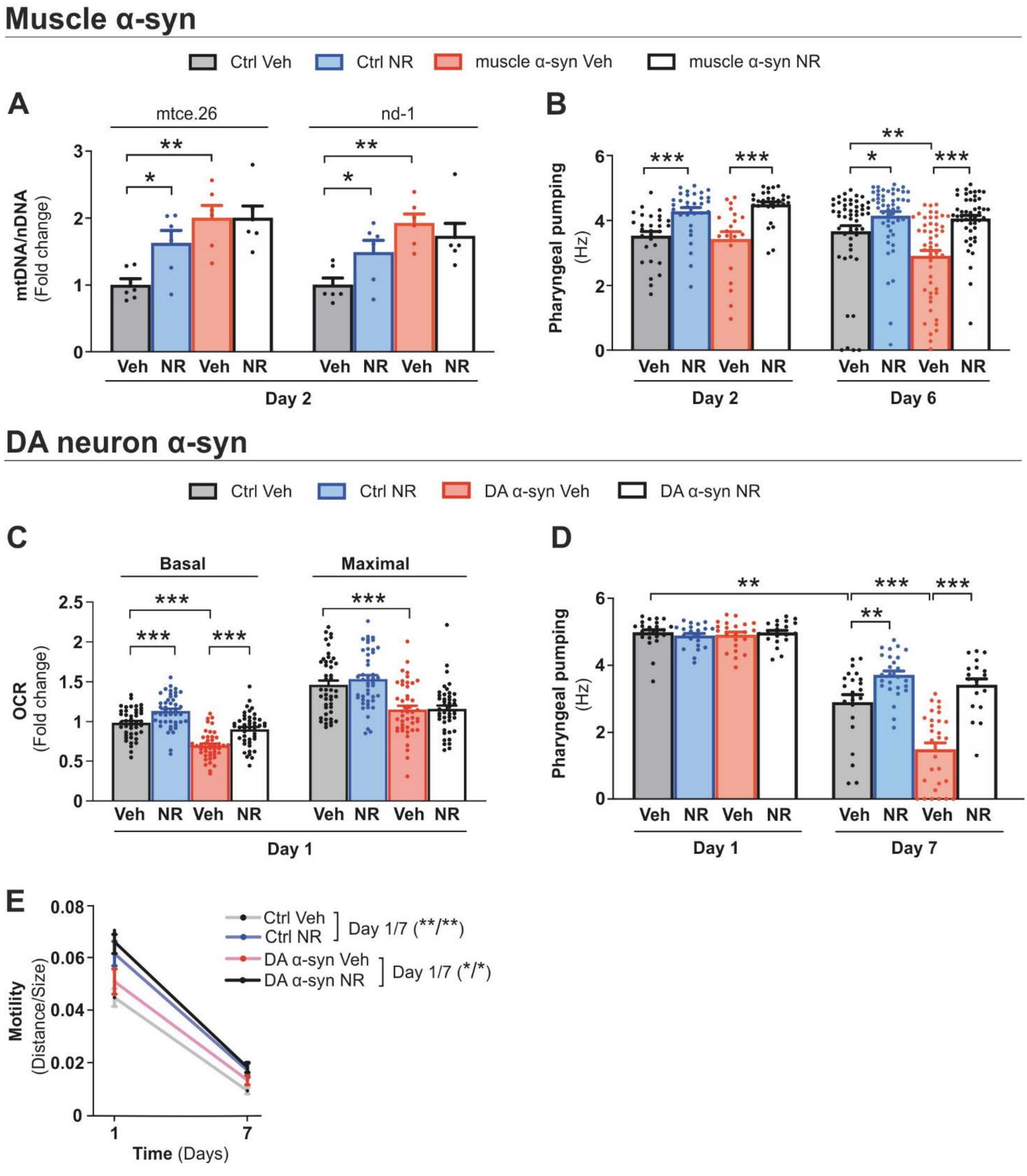


**Supplemental Figure 1. NR supplementation improves food transport and mitochondrial health in *C. elegans* strains overexpressing a-synuclein.**

(**A**) Relative mtDNA amount in the control and muscle α-syn strains after vehicle and NR supplementation at day 2 of adult age (n=6 biologically independent samples from 2 independent experiments). (**B**) Quantification of pharyngeal pumping activity as a measure of food transport in the control and muscle α-syn strains after vehicle and NR supplementation at days 2 and 6 of adult age. Each data point represents the mean of 2 independent experiments with 20 to 50 animals per group.

(**C**) Basal and maximal mitochondrial respiration measured in the control and DA α-syn strains after vehicle and NR supplementation at day 1 of adult age. Each data point represents the mean of 3 independent experiments in total with 45 wells per group, each group having 5 to 8 animals per well. (**D)** Quantification of pharyngeal pumping activity in the control and DA α-syn strains after vehicle and NR supplementation at days 1 and 7 of adult age. Each data point represents the mean of 2 independent experiments with 19 to 29 animals per group. (**E)** Motility (distance travelled normalized to the size of the animals) in the control and DA α-syn strains with vehicle or NR supplementation at days 1 and 7 of adult age. Each data point represents the mean of 2 independent experiments with 36 to 92 animals per group.

All data are mean ± SEM. *$P < 0.05$; **$P \leq 0.01$; ***$P \leq 0.001$; ns, not significant. Statistical analysis was performed in GraphPad prism version 9.0.0. Overall differences between conditions were assessed in one-way ANOVA using an uncorrected version of Fisher's LSD test.

### SUPPLEMENTAL FIGURE 2

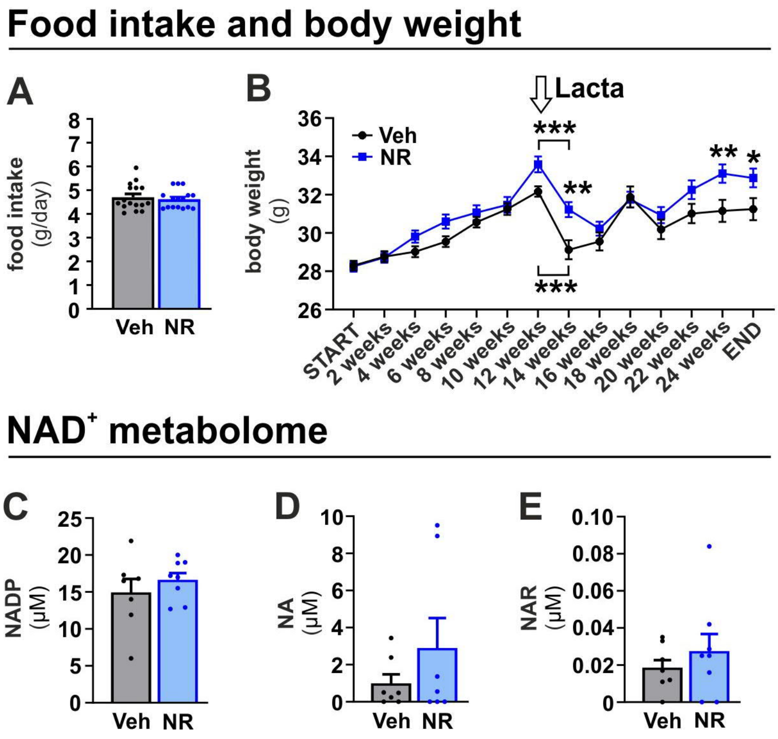


**Supplemental Figure 2. NR does not alter food intake or the levels of NADP, NA and NAR but improves overall well-being of mice after lactacystin injection.**

(**A**) Food intake in mice fed with a vehicle (Veh) or NR-enriched diet (NR). The measurement was performed twice: at 20 weeks and 22 weeks after the experiment was started. Results are expressed as an average of the two independent measurements. Veh (n = 16); NR (n = 16). (**B**) Weight development in mice that received vehicle (Veh) or NR-enriched diet (NR). Lactacystin was injected 12 weeks after NR treatment started. Veh (n = 38); NR (n = 38) before lactacystin injection; Veh (n = 34); NR (n = 38) after lactacystin injection. Cerebellar content of (**C**) NADP; (**D**) NA, and (**E**) NAR in vehicle and NR supplemented mice at the 12-week time point. Veh (n = 7) and NR (n = 8). Lacta, lactacystin; NA, nicotinic acid; NADP, nicotinamide adenine dinucleotide phosphate; NAR, nicotinic acid riboside; NR, nicotinamide riboside; Veh, vehicle.

Data are presented as mean ± SEM. $^{**}P \leq 0.01$, $^{***}P \leq 0.001$. Statistical analysis was performed using GraphPad Prism version 8.0.2. Overall differences were assessed with Student's t-test followed by Welch's correction (**A**), two-way repeated measures ANOVA followed by Sidak's multiple comparison test (**B**), and with one-way ANOVA followed by an uncorrected version of Fisher's LSD test (**C-E**).

### SUPPLEMENTAL FIGURE 3

#### VTA, Lacta 12w

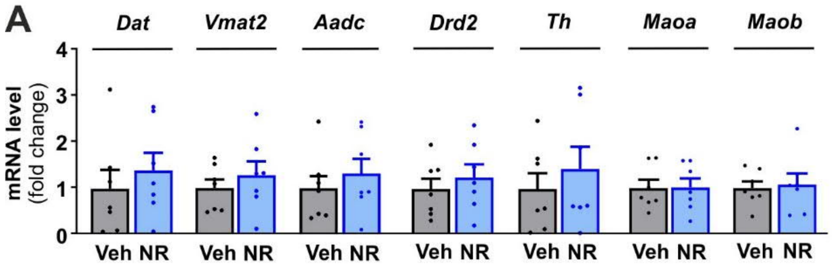


#### SN, Lacta 12w

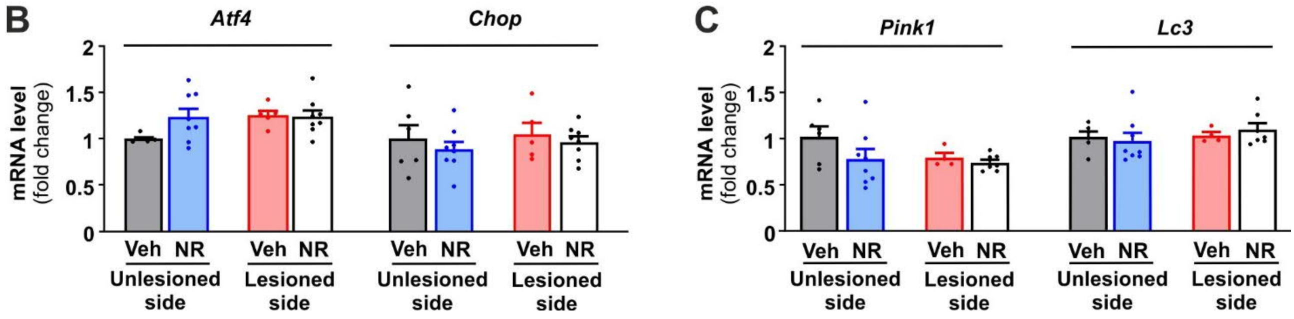


**Supplemental Figure 3. Long-term NR supplementation results in downregulation of DA metabolism specifically in the substantia nigra upon proteostasis failure caused by lactacystin.**

(**A**) Gene expression of *Dat*, *Vmat2*, *Aadc*, *Drd2*, *Th*, *Maoa*, and *Maob* mRNA quantified by qRT-PCR in the ventral tegmental area of vehicle and NR supplemented mice at 12 weeks after lactacystin injection. Veh (n = 7); NR (n = 7). (**B**) Gene expression level of relevant genes involved in UPR$^{er}$ signalling and (**C**) mitochondrial stress defense and autophagy quantified by qRT-PCR in the substantia nigra of vehicle and NR supplemented unlesioned and lactacystin-lesioned mice 12 weeks after lactacystin injection. Veh, unlesioned side (n = 6); NR, unlesioned side (n = 8); Veh, lesioned side (n = 5); NR, lesioned side (n = 8).

Data are presented as mean ± SEM. Statistical analysis was performed using GraphPad Prism version 8.0.2. Overall differences between conditions were assessed with one-way ANOVA followed by an uncorrected version of Fisher's LSD test. Aadc, aromatic L-amino acid decarboxylase; Atf4, activating transcription factor 4; Chop, C/EBP homologous protein; Dat, dopamine active transporter; Drd2, dopamine receptor 2; Lacta, lactacystin; Lc3, microtubule-associated protein 1A/1B-light chain 3; Maoa, monoamine oxidase A; Maob, monoamine oxidase B; Pink1, PTEN-induced kinase 1; NR, nicotinamide riboside; SN, substantia nigra; Th, tyrosine hydroxylase; Veh, vehicle; Vmat2, vesicular monoamine transporter 2; VTA, ventral tegmental area.

### SUPPLEMENTAL FIGURE 4

#### Study design, No Lacta

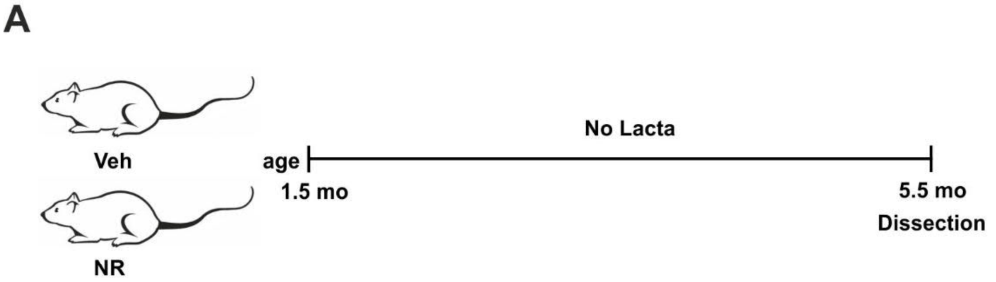


#### SN, No Lacta

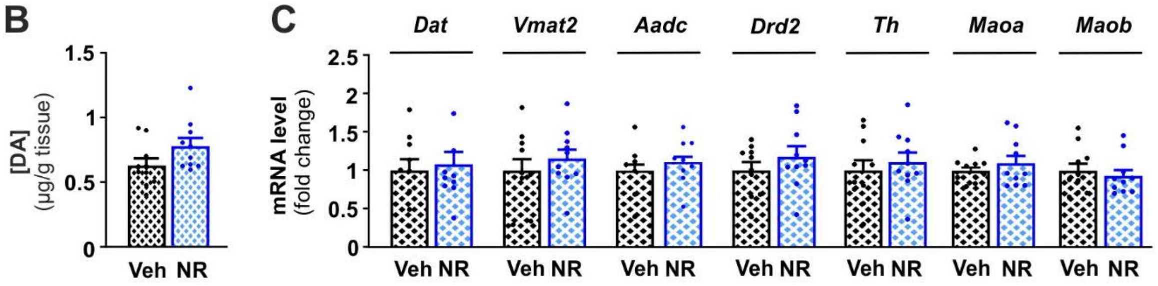


**Supplemental Figure 4. NR has no effect on DA levels and DA metabolism in the substantia nigra of mice that did not undergo lactacystin injection.**

(**A**) Experimental design for mice who received either a vehicle (Veh) or NR-enriched diet (NR) and were not subjected to lactacystin injections (No Lacta). (**B**) DA tissue level and (**C**) gene expression of *Dat*, *Vmat2*, *Aadc*, *Drd2*, *Th*, *Maoa*, and *Maob* in the substantia nigra of vehicle and NR supplemented uninjected mice. Veh (n = 10); NR (n = 10).

Data are presented as mean ± SEM. Statistical analysis was performed using GraphPad Prism version 8.0.2. Overall differences between conditions were assessed with one-way ANOVA followed by an uncorrected version of Fisher’s LSD test. Aadc, aromatic L-amino acid decarboxylase; DA, dopamine; Dat, dopamine active transporter; Drd2, dopamine receptor 2; Lacta, lactacystin; Maoa, monoamine oxidase A; Maob, monoamine oxidase B; NR, nicotinamide riboside; SN, substantia nigra; Th, tyrosine hydroxylase; Veh, vehicle; Vmat2, vesicular monoamine transporter 2.

**Supplemental Table 1. List of primers used in *C.elegans*.**

| | | |
|---|---|---|
| *hsp6* | Forward | AGAGCCAAGTTCGAGCAGAT |
| | Reverse | TCTTGAACAGTGGCTTGCAC |
| *hsp60* | Forward | GGAAGCCCAAAGATCACAAA |
| | Reverse | CAGCCTCCTCATTAGCCTTG |
| *ymel1* | Forward | CAAAACCTGATCTCGCTGGG |
| | Reverse | TTCTCAATGTCGGCTCCAGT |
| *lonp1* | Forward | CGATGATGGCCATTGTGCAG |
| | Reverse | CGCTTTGAAACATCAATTTCATCCA |
| *clpp-1* | Forward | TGATAAGTGCACCAGTGTCCA |
| | Reverse | TGATTCTGGAGTTCGGGAGA |
| *ubl-5* | Forward | TGATCGCTGCACAAACTGGAACAC |
| | Reverse | CCCTCGTGAATCTCGTAATCCATC |
| *Y45F10D.4* | Forward | CGAGAACCCGCGAAATGTCGGA |
| | Reverse | CGGTTGCCAGGGAAGATGAGGC |
| *cdc-42* | Forward | AGGAACGTCTTCCTTGTCTCC |
| | Reverse | GGACATAGAAAGAAAAACACAGTCAC |
| *mtce.26* | Forward | GGTTGTGGGACTAGGTGAACA |
| | Reverse | CAGGGTGCCCCATTGTTCTT |
| *nd-1* | Forward | AGCGTCATTTATTGGGAAGAAGAC |
| | Reverse | AAGCTTGTGCTAATCCCATAAATGT |
| *act-3* | Forward | TGCGACATTGATATCCGTAAGG |
| | Reverse | GGTGGTTCTCCGGAAAGAA |

**Supplemental Table 2. List of primers used in mice.**

| | | |
|---|---|---|
| *Dat* | Forward | AACCTGTACTGGCGGCTATG |
| | Reverse | GCTGACCACGACCACTACA |
| *Vmat2* | Forward | ATGCTGCTCACCGTCGTAGT |
| | Reverse | TTTTTCTCGTGCTTAATGCTGT |
| *Aadc* | Forward | GCATTGAGGGTCGTCCAGTG |
| | Reverse | CTGGGTATGAGCTAGCCGTG |
| *Drd2* | Forward | ACACACGCTACAGCTCCAAG |
| | Reverse | GGAGTAGACCACGAAGGCAG |
| *Th* | Forward | CCCAAGGGCTTCAGAAGAG |
| | Reverse | GGGCATCCTCGATGAGACT |
| *Maob* | Forward | GCCCAGTATCACAGGCCAC |
| | Reverse | CGGGCTTCCAGAACCAAGA |
| *Maob* | Forward | GCACTGAAACAGCCTCACAC |
| | Reverse | TCGTGCAGGGACATCCAAAG |
| *Atf4* | Forward | CTCATGGGTTCTCCAGCGACAAG |
| | Reverse | GTCAAGAGCTCATCTGGCATGG |
| *Chop* | Forward | GAAGCCTGGTATGAGGATCTGCAGGAGGTC |
| | Reverse | CTTTGGGATGTGCGTGTGACCTCTGTTG |
| *Pink1* | Forward | GCTTGCCAATCCCTTCTATG |
| | Reverse | CCCCACCAAGATCCCAGT |
| *Lc3* | Forward | CTCTCGCTGGAGCAGTGAC |
| | Reverse | CGCTCATGTTCACGTGGT |
| *Rps18* | Forward | GATGGGAAGTACAGCCAGGTTC |
| | Reverse | AAAGTGGCGCAGCCCTCTA |
| *Gapdh* | Forward | CCTCGTCCCGTAGACAAAA |
| | Reverse | ATGAAGGGGTCGTTGATGGC |
| *Actb* | Forward | CTGTCGAGTCGCGTCCA |
| | Reverse | ACGATGGAGGGGAATACAGC |
